# Resurrection of ancestral effector caspases identifies novel networks for evolution of substrate specificity

**DOI:** 10.1101/749143

**Authors:** Robert D. Grinshpon, Suman Shrestha, James Titus-McQuillan, Paul T. Hamilton, Paul D. Swartz, A. Clay Clark

## Abstract

Apoptotic caspases evolved with metazoans more than 950 million years ago (MYA), and a series of gene duplications resulted in two subfamilies consisting of initiator and effector caspases. The effector caspase genes (caspases-3, -6, and -7) were subsequently fixed into the Chordata phylum more than 650 MYA when the gene for a common ancestor (CA) duplicated, and the three effector caspases have persisted throughout mammalian evolution. All caspases require an aspartate residue at the P1 position of substrates, so each caspase evolved discrete cellular roles through changes in substrate recognition at the P4 position combined with allosteric regulation. We examined the evolution of substrate specificity in caspase-6, which prefers valine at the P4 residue, compared to caspases-3 and -7, which prefer aspartate, by reconstructing the CA of effector caspases (AncCP-Ef1) and the CA of caspase-6 (AncCP-6An). We show that AncCP-Ef1 is a promiscuous enzyme with little distinction between Asp, Val, or Leu at P4. The specificity of caspase-6 was defined early in its evolution, where AncCP-6An demonstrates preference for Val over Asp at P4. Structures of AncCP-Ef1 and of AncCP-6An show a network of charged amino acids near the S4 pocket that, when combined with repositioning a flexible active site loop, resulted in a more hydrophobic binding pocket in AncCP-6An. The ancestral protein reconstructions show that the caspase-hemoglobinase fold has been conserved for over 650 million years and that only three substitutions in the scaffold are necessary to shift substrate selection toward Val over Asp.

## Introduction

The caspase family of proteases offers an attractive model for examining protein evolution because a common protein scaffold (the caspase-hemoglobinase fold) was used to develop subfamilies that differ in oligomeric states, enzyme specificity, and allosteric regulation. Caspase genes predate multicellularity [1] and are represented in all kingdoms of life [2]. Caspase proteases are thought to have evolved from an ancestral immune system into two general classes, inflammatory or apoptotic caspases, consisting of over twelve proteins [3, 4]. Within the apoptotic caspases, two subfamilies further evolved into apoptotic initiators or effectors [3, 5]. Following additional gene duplications, an ancestral initiator caspase gave rise to four genes (caspases-8,-10,-18, c-FLIP), while an ancestral effector caspase gave rise to three genes (caspases-3,-6,-7) (Figure 1A) [6].

**Figure 1.**
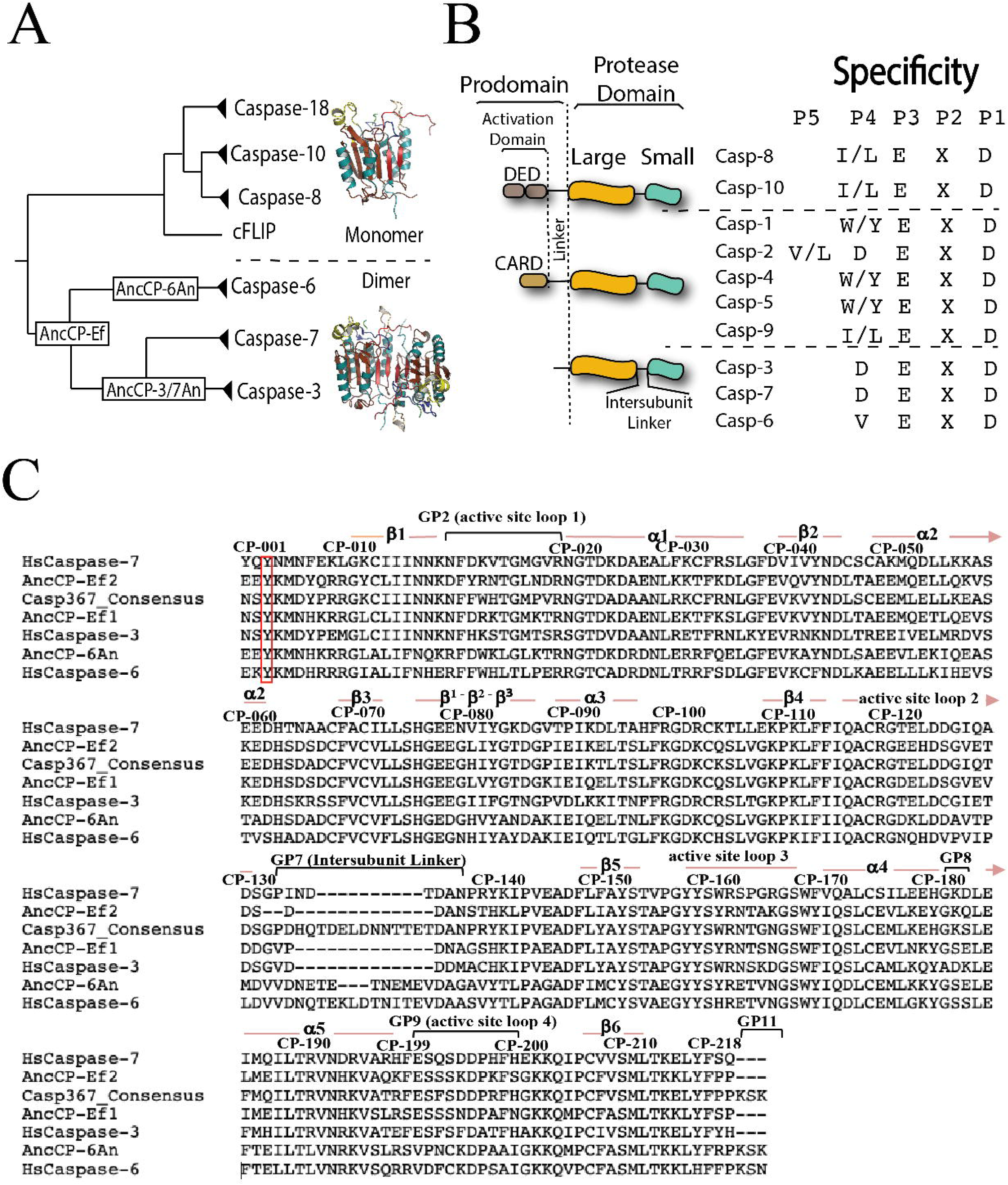
Ancestral caspase proteins. (A) Evolutionary relationship of initiator (monomer) and effector (dimer) caspases. AncCP-Ef is the common ancestor to effector caspases-3, -6, and -7. AncCP-6An is the common ancestor to the caspase-6 lineage. (B) Domain organization and enzyme specificity for ten human caspases. DED refers to death effector domain, and CARD refers to caspase activation and recruitment domain. The caspase protomer consists of a large and small subunit separated by an intersubunit linker (IL), which is cleaved upon maturation, and the active caspase is a dimer of protomers. All caspases recognize aspartate at the P1 position of the substrate, and specficity is primarily determined by the amino acid at the P4 position. (C) Sequences of human (Hs) caspases-3, -6, and -7 compared to two reconstructions of AncCP-Ef (AncCP-Ef1, -Ef2) and AncCP-6An. The Caspase-367 concensus sequence was determined from 667 sequences of extant effector caspases in the CaspBase. Numbering refers to the common position, as described in the text, and secondary structural elements and active site loops are indicated. Red box indicates CP-Y001, which identifies the start of the protease domain.

Caspases are produced in the cell as inactive zymogens, and the oligomeric form of the zymogen is a key to regulation [7–9]. The initiator caspases are monomers, and the ability to form heterodimers *versus* homodimers in response to cellular conditions is a key characteristic in cell fate decisions regarding the activation of necroptosis or apoptosis pathways [10, 11]. In contrast, effector caspases evolved as obligate homodimers that are processed by initiator caspases, so discrete cellular functions of effector caspases developed through a combination of changes in substrate specificity and allosteric regulation [9, 12]. Studies of enzyme families have identified features that contribute to enzyme specificity [13–15], but components that determine specificity of the caspase active sites have been elusive. Enzyme active sites provide the stereo-selective environment for reaction ground or transition states, or the protein scaffold provides the proper conformational dynamics that facilitate substrate binding and reaction chemistry [16, 17], so there may be multiple combinations of residues that provide the proper environment within the context of the protein scaffold [18, 19].

Caspases cleave target proteins through recognition of a tetrapeptide motif with the noted exception of caspase-2, which recognizes a pentapeptide sequence [20]. In some cases, enzyme specificity is coupled to exosites that facilitate substrate selection [21–25]. Positions P1-P4 on the peptide are coordinated by their corresponding substrate pocket, S1-S4 in the active-site, and the P1 residue is almost always an aspartate [26]. Because specificity is determined primarily by the amino acid at the P4 position, caspases are sub-categorized into three groups based on recognition for the P4 amino acid: group I prefers a bulky residue (W,H); group II prefers hydrophilic residues (D,E); and group III prefers aliphatic residues (I,L,V) (Figure 1B). Although the effector caspases are relatively closely related, caspases-3 and -7 are characterized as group II specificity, while caspase-6 shows group III specificity. The selection at P4 (D vs V) results in overlapping but nonidentical substrate profiles based on degradome analyses [27–30]. In the evolution of chordates, new caspase substrate specificities were important in developmental stages of the brain and nervous systems [31–34], so what may appear to be subtle changes in enzyme selection have large consequences in cellular development.

Current models suggest that modern enzymes evolved from promiscuous ancestral proteins through amino acid substitutions that were coupled to the selection of pre-existing suboptimal activity [35–37]. While the caspase-8 subfamily evolved into cell fate determinants, with largely uniform substrate selection ((I/L)EXD), changes in effector caspases resulted in two distinct specificities-DxxD *versus* VxxD. The evolutionary trajectories that resulted in the distinct substrate specificities are not known. Although horizontal studies that compare extant enzymes may identify the importance of key active site residues, they rarely uncover the set of residues that are responsible for functional diversity in large protein families [38]. Generally, substitutions from horizontal studies lack evolutionary context, where protein epistasis affects the specific combination of amino acids along discrete evolutionary paths [39, 40]. Directed evolutionary approaches expand the sequence space that can be examined, and such studies identified a combination of amino acids that relax specificity in caspase-7, resulting in a shift in substrate cleavage profiles *in cellulo* of evolved-caspase-7 enzymes [30]. Evolutionary biochemical methods further expand the sequence space to include the entire protein, yet the methods simultaneously narrow the scope of the problem by also examining changes that occurred between common ancestral proteins [41].

To determine the evolutionary pathways leading to VxxD *versus* DxxD specificity in effector caspases, we reconstructed sequences and resurrected ancestral proteins for the common ancestor (CA) of caspases-3, -6, and -7 (called AncCP-Ef1) and for the CA of the caspase-6 branch (AncCP-6An), where “An” refers to anamniotes. We show that AncCP-Ef1 is indeed a promiscuous enzyme that exhibits low activity and little preference for Val, Asp, or Leu at the P4 position. The selection of Val over Asp occurred early in the evolution of caspase-6, with AncCP-6An. Structures of AncCP-Ef1 and of AncCP-6An determined by X-ray crystallography show a unique mechanism of introducing a network of charged amino acids that increase hydrophobicity of the S4 binding pocket in the caspase-6 evolutionary pathway. Introduction of the network into AncCP-Ef1 shifted specificity to Val over Asp, as in caspase-6. Together, the data demonstrate that only three amino acid substitutions are required in the ancestral scaffold to shift specificity in the caspase-6 proteins.

## Materials and methods

### Ancestral protein reconstruction

Two lists of taxa were used for ancestral protein reconstruction (APR). One list (APR_1, Supplemental Information, Table S1) was generated using a precursor to the CaspBase [42] (caspbase.org), and one list (APR_2, Supplemental Information, Table S2) queried the CaspBase. Each list has high sequence coverage spanning the majority of known proteins within the caspase family. While APR_1 has a total of 253 caspase sequences, APR_2 has a total of 258 caspase sequences. APR_2 emphasizes mammal lineages over non-mammal lineages, with 127 mammal caspase sequences, while APR_1 has 82 mammalian caspase sequences. Care was taken in both lists to mitigate erroneous sequences, to eliminate incomplete lineages sorting by including high coverage across all representative taxa from all major vertebrate groups, and to incorporate full gene tree representation of each known caspase family member within each vertebrate group. The prodomain was pruned from our sequences because the prodomains have high sequence variations and lengths in the caspase family, and their inclusion results in missing data and noise to downstream analyses. The multiple sequence alignment (MSA) was computed using PROMALS3D [43, 44]. Alignments were checked in Geneious [45] to assess alignment accuracy, and we utilized Prottest 3 [46] to generate the proper model for phylogenetic analysis using AICc (Akaike Information Criterion) weights to gather the highest probable model of protein evolution [47]. The phylogenetic tree was generated using IQTREE [48], using a combination of hill-climbing approaches and stochastic perturbation methods for accuracy and time-efficiency, and the tree was bootstrapped 1000 times as a test of phylogeny [49]. The tree was examined to remove erroneous sequences, mislabels, and to mitigate missing data, resulting in a highly effective alignment for APR. The APRs were constructed with FastML [50], using codon-bases reconstruction models for accuracy since the models were generated from whole annotated genomes with complete metadata. We used a LG model of substitution [51] generated by Prottest 3, and our framework used maximum likelihood (ML) for indel reconstruction. We provided our ML tree as a guide, optimizing branch lengths with highly divergent sequences, set gamma distribution, and computed a joint reconstruction to generate APRs at each node of interest. Sequences were codon-optimized for expression in *E. coli*, cloned into pET11a vector and included a C-terminal His6-tag (GenScript, USA). The AncCP-6An was also designed similarly to the caspase-6 CT (constitutive two-chain) construct described previously [52]. All proteins were purified as described previously [53–55].

### Crystallization and data collection

Each protein was dialyzed in a buffer of 10 mM Tris-HCl, pH 8.5, and 1 mM DTT, concentrated to 8-10 mg/mL, and stored at -80°C. The molar extinction coefficients for the APRs were determined by ProtParam under reduced conditions [56] (Supplemental Information, Table S3). The inhibitors (Ac-DEVD-CHO (acetyl-Asp-Glu-Val-Asp-aldehyde) or Ac-VEID-CHO (acetyl-Val-Glu-Ile-Asp-aldehyde) in DMSO) were added at 1:5 (w/w) ratio of protein:inhibitor, and solutions were incubated on ice for 1 hour in the dark. Initial crystallization conditions were found using Hampton crystal screens (crystal screen 1 and PEG/ion screen 1). For each well, a solution of the screen (490 μL), DTT (5 μL of 1 M solution), and sodium azide (5 μL of 300 mM solution) were added. Crystals grew using the hanging drop vapor diffusion method at 18°C using 4 μL drops that contained equal volumes of protein and reservoir solutions. For AncCP-Ef1(DEVD), optimal conditions were found in a solution of 0.2 M ammonium acetate, pH 4.6, 0.1 M sodium acetate trihydrate, 30% PEG 4000. For AncCP-6An(VEID), optimal conditions were found in a solution of 0.2 M ammonium fluoride, pH 6.2, 20% PEG 3350. Crystals were flash frozen in liquid nitrogen following the addition of 20% MPD (2-methylpentane-2,4-diol) or 20% glycerol plus well buffer. Data were collected at 100K at the SER-CAT synchrotron beamline (Advance Photon Source, Argonne National Laboratory Argonne, IL, U.S.A.). Each data set contained 180 frames at 1^°^ rotation. The proteins crystallized in the orthorhombic space group P2_1_2_1_2_1_ and were phased with a previously published human CASP3 structure (PDB entry 2J30). Data reduction and model refinements were done using HKL2000, COOT, and Phenix, and a summary of the data collection and refinement statistics is shown in Supplemental Information, Table S4.

### Substrate-phage display

Enzyme specificity was determined by substrate-phage display as described previously [57]. Briefly, phage libraries consisting of caspase recognition sequences, with either random or fixed (aspartate) P1 position, were bound to Ni-NTA resin. Enzyme (10-100 nM) was added to initiate the reaction, and samples were incubated between 1 and 20 hours. *E. coli* ER2738 cells were used to amplify the previous round by infecting cells with the supernatant from the reaction. The cells were grown for 4 hours, and the supernatant was collected and used as the library for the following round of selection. Colony counting was used to determine the endpoint of the experiment, when the amount of library bound to the resin was similar to the amount released during the treatment. The amount of phage released during the reaction *versus* the control (without enzyme) was monitored to ensure progress in substrate selectivity.

### Enzyme activity assay

Enzyme activity was determined in a buffer of 150 mM Tris-HCl, pH 7.5, 50 mM NaCl, 10 mM DTT, 1% sucrose, 0.1% CHAPS (assay buffer) at 25 °C, as previously described [58, 59]. The total reaction volume was 200 µL, and the final enzyme concentration was 10 nM. Following the addition of substrate (Ac-DEVD-AFC (acetyl-Asp-Glu-Val-Asp-7-amino-4-trifluoromethylcoumarin), Ac-LETD-AFC (Leu-Glu-Thr-Asp-7-amino-4-trifluoromethylcoumarin), or Ac-VEID-AFC (acetyl-Val-Glu-Ile-Asp-7-amino-4-trifluoromethylcoumarin), the samples were excited at 400 nm, and emission was monitored at 505 nm for 60 seconds. The steady-state parameters, K_M_ and k_cat_, were determined from plots of initial velocity *versus* substrate concentration and are presented in Supplemental Information, Table S5.

### Data Deposition

Crystal structures have been deposited in the Protein Data Bank, www.wwpdb.org (PDB ID codes: 6PDQ, AncCP-Ef1(DEVD); 6PPM, AncCP-6An)

## Results

### Reconstruction of ancestral effector caspases

A comparison of human caspases-3 and -6 shows that there are 118 differences out of 237 amino acids, not including the intersubunit linker or prodomain (Figure 1C). Many of the differences (Figure 2A, orange) map to three active site loops (called L1, L3, and L4) as well as the five α-helices on the surface of the protein. In particular, helix 2 (H2) and the adjoining short surface strands (β^1^-β^3^) are known to undergo a coil-to-helix transition in caspase-6, which allosterically inhibits the enzyme [60]. Likewise, in caspase-3 a conserved network of water molecules binds to helices 1, 4, and 5 to couple an allosteric site in the dimer interface to changes in the active site [61]. While each of the 118 sites that differ between caspases-3 and -6 could be considered for horizontal mutations, we reasoned that differences near the active site were more likely to affect substrate specificity while changes in the surface helices may be coupled to allosteric regulation. In addition, rather than simply swapping activity between the two enzymes, we wanted to determine the evolutionary changes in caspases that resulted in specificity of the extant enzymes.

**Figure 2.**
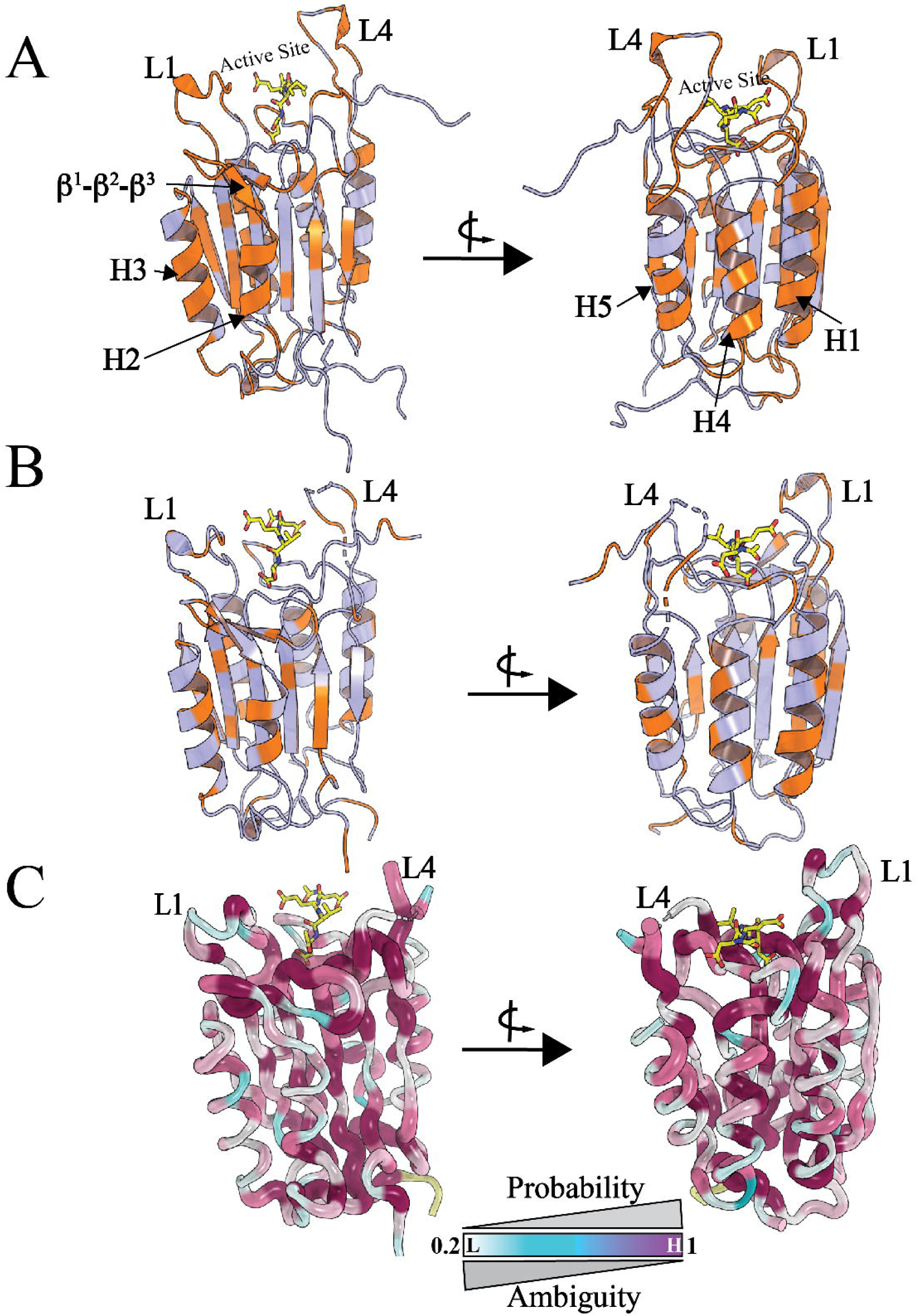
Comparison of changes in extant and ancestral caspases. (A) Site-specific differences between HsCaspase-3 and HsCaspase-6 are shown in orange, mapped onto the HsCaspase-6 protomer (PDB ID: 3OD5). The five surface helices (H1-H5), short surface β-strands (β^1^-β^3^) and two of the five active site loops (L1, L4) are indicated. (B) Site-specific differences (orange) between AncCP-Ef1 and AncCP-6An. (C) Site-specfic posterior probabilities for AncCP-Ef. In the putty representation, regions of higher probability are shown as thicker lines.

In order to examine changes that occurred in the active site binding pockets, we first queried the CaspBase [42] for effector caspase sequences, and based on the 667 sequences returned, we generated a consensus sequence for effector caspases (Figure 1C). We previously developed the common position (CP) numbering scheme in order to compare positions in evolutionarily divergent caspases [42], where the CP system describes common positions in all caspases as well as more divergent regions, such as active site loops (called gapped positions, or GP). Here, the amino acid position for each caspase is shown by superscript, whereas the common position is preceded by “CP-.” The active site S1 binding pocket of caspase-6 is the same as all other caspases and consists of Arg^64^ (CP-018), Q^161^ (CP-115), and Arg^220^ (CP-161) (Supplemental Information, Figure S1). The S2 binding pocket is formed by the side chains of Tyr^217^ (CP-158), His^168^ (CP-122), and His^219^ (CP-160). While Tyr^217^ (CP-158) is highly conserved in effector caspases, CP-122 and CP-160 are both highly conserved (>90%) histidine residues that are unique to caspase-6. The S3 binding pocket is partially formed with Arg^220^ (CP-161), and the S4 binding pocket is formed by His^219^ (CP-160), Glu^221^ (CP-162), Trp^227^ (CP-168), and Val^261^ (GP9-V01) [62]. In caspase-6, CP-162 is conserved as glutamate, whereas caspases with group II substrate specificity (caspase-2, -3, and -7) utilize asparagine (Figure 1C). In addition, GP9-01 is conserved as valine in caspase-6, whereas the group II caspases typically utilize a glutamate that is not conserved.

To determine the evolutionary changes in the active site that led to modern substrate selection, we reconstructed common ancestral proteins for the effector caspases and for caspase-6 (Figure 1A). Ancestral protein reconstruction (APR) techniques can reveal amino acid substitutions that result in neofunctionalization of proteins by creating an evolutionary map leading to the new functions [63]. The effector caspases diverged from a common ancestor approximately 650 million years ago into the caspase-6 and caspase-3/-7 branches, and caspases-3 and -7 diverged into separate branches later (Figure 1A). Utilizing data from the CaspBase, we reconstructed proteins at the node that represents the common ancestor of caspase-3/6/7, called AncCP-Ef1, and for the CA of caspase-6, called AncCP-6An (Figure 1C). The data show only 42 sites changed from AncCP-Ef1 to AncCP-6An, referred to as vertical substitutions, with few changes in active site loops L1 and L4 (Figs. 1C and 2B).

APR analyses result in site-specific probabilities for each position in the protein, with the ultimate goal to examine the characteristics of the protein rather than the precise ancestral sequence [63, 64]. Thus, the predictions made with APR may differ for each input dataset such that the nodes of a phylogenetic tree actually represent a pool of possible ancestors. Due to evolutionary divergence for each site, the APR analysis identifies sites that may be ambiguous, defined here as a site that has less than 70% probability. For the caspase APRs, ambiguous sites generally occurred in two types, those in which two amino acids show nearly equal probability (∼50% each) in leading to extant proteins (here referred to as A1 or ambiguity type 1), or those in which multiple amino acids show lower probabilities (here referred to as A2 or ambiguity type 2). In the first case, A1, the amino acids are generally conserved, but differ, for two branches of a family. In the second case, A2, the sites are generally less conserved among family members. For AncCP-Ef1, we mapped the site-specific probability onto the protein structure (Figure 2C), and the data show low ambiguity in the active site and protein core, and higher ambiguity in the surface helices, particularly helices 2 and 3. The data are similar to conservation maps of extant enzymes, which display lower conservation in the surface helices [23]. Together, the data suggest that the ambiguous sites are largely due to highly variant regions in extant caspases.

To examine the robustness of the AncCP-Ef reconstruction, we carried out two separate APR experiments by using different datasets and comparing their posterior probabilities. The two proteins, called AncCP-Ef1 and AncCP-Ef2, represent the same pool of possible ancestors from the different reconstructions, and both proteins were resurrected to corroborate the experimental results. For cloning purposes, we used the prodomain sequences from caspases-3 or -6 from *Homo sapiens* and intersubunit linker (IL) sequences computed from the APRs (Figure 1C). Previous studies showed that the prodomain does not affect enzymatic function *in vitro* [65], and the sequences are removed during zymogen maturation. In addition, we estimated the length of the IL as the average length in the organisms that diverged around the time of the ancestral node, and the reconstructed residues were used for the IL of that node. For example, AncCP-6An represents the ancestor of caspase-6 up to bony fish, so we used the average length of the IL for Actinopterygii (ray-finned fishes) for reconstructing AncCP-6An. In this case, the IL is three residues shorter than that of human caspase-6 (Figure 1C).

A comparison of the protein sequences shows that while human caspases-3 and -6 are only 40% identical, AncCP-Ef1 and AncCP-6An have higher sequence identity with each other (68%) and with extant enzymes (54-75%) (Supplemental Information, Figure S2). In addition, the data show that the ancestral enzymes are more acidic compared to extant enzymes, with calculated pI of 5.2-5.4 *versus* 5.7-6.5, due to higher percentages of glutamate and aspartate (Supplemental Information, Table S3). The two reconstructions from the same ancestral node, AncCP-Ef1 and AncCP-Ef2, have 81% sequence identity and differ in 39 sites, not including the IL (Figure 1C, Supplemental Information, Figure S3). While several sites in helices 1 and 4 vary, many of the differences between AncCP-Ef1 and AncCP-Ef2 occur in active site loops L1 and L4. For example, in AncCP-Ef1 the sequence (CP-195)VSLRS(CP-199) is more similar to that of human caspase-6 (VSQRR) than that of human caspase-3 (VATEF) (Figure 1C), while the opposite is true for AncCP-Ef2 (VAQKF). In both reconstructions, however, the amino acid at position GP9-01, which forms part of the S4 binding pocket, is well-determined as glutamate. As described below, the differences between AncCP-Ef1 and AncCP-Ef2 have little effect on enzyme activity.

### Ancestral effector caspase was a promiscuous enzyme

We examined the enzyme activity of AncCP-Ef1 and of AncCP-Ef2 against three tetrapeptide sequences (DEVD, VEID, and LETD), representing specificities for caspases-3, -6, and -8, respectively (Supplemental Information, Table S5). As described previously, k_cat_/K_M_ values are in the range of ∼0.5-2×10^5^ M^-1^s^-1^ for human caspases on the optimal P4 residue [20,26,66], with k_cat_ values generally ∼0.3 to 1 s^-1^ and K_M_ values of ∼15 μM. In the case of AncCP-Ef1, k_cat_ values were similar to those of extant caspases, but the activity was >5-fold lower due to significantly higher K_M_ values (>90 μM), resulting in activities of ∼10^3^ M^-1^s^-1^. A comparison of the two ancestral effector caspase reconstructions shows that AncCP-Ef1 and AncCP-Ef2 exhibited similar activity for DEVD substrate (2.8-6.8×10^3^ M^-1^s^-1^), but the activity of AncCP-EF2 for VEID was below the detection limit for the assay (∼5×10^2^ M^-1^s^-1^) due to high K_M_ values (Supplemental Information, Table S5). Together, the data for AncCP-Ef1 and AncCP-Ef2 show that the CA of effector caspases has low activity for the three substrates, due to high K_M_ values, indicating weaker substrate binding compared to extant enzymes. There was a significant improvement in the activity of AncCP-6An compared to the common ancestor, particularly regarding valine at P4 (Supplemental Information, Table S5). In this case, we find ∼3-fold increase in k_cat_/K_M_ with Val, little change with P4 Leu, and no activity against Asp at P4. The change in activity for AncCP-6An was due to a decrease in K_M_.

We also compared the substrate specificity of AncCP-Ef1 and of AncCP-6An using substrate-phage selection. As described previously [57], we utilized two substrate-phage libraries, either with randomized P5-P1’ positions or with aspartate fixed at the P1 position, and the data were combined to define substrate selection. As shown in Figure 3A, HsCaspase-6 shows preference (P5-P1’) for (L/S)(T/V)EVDA. In contrast, AncCP-Ef1 was less selective for the P4 amino acid, with Leu, Ile, Asp being the most prevalent (Figure 3B) and an overall P5-P1’ selection of T(L/I/D)E(T/V)DG. In contrast, AncCP-6An showed a preference for Val at P4, with an overall P5-P1’ selection of Y(V/T)LTDS (Figure 3C), which is consistent with the change in specificity as shown in the tetrapeptide activity studies. Together, the enzyme activity data on small peptides and on substrate-phage libraries show that the ancestral effector caspase had >10-fold lower activity compared to extant enzymes, and with little preference for Asp, Val, or Leu. In contrast, the specificity of caspase-6 for hydrophobic *versus* charged amino acids at P4 arose in the early caspase-6 ancestor.

**Figure 3.**
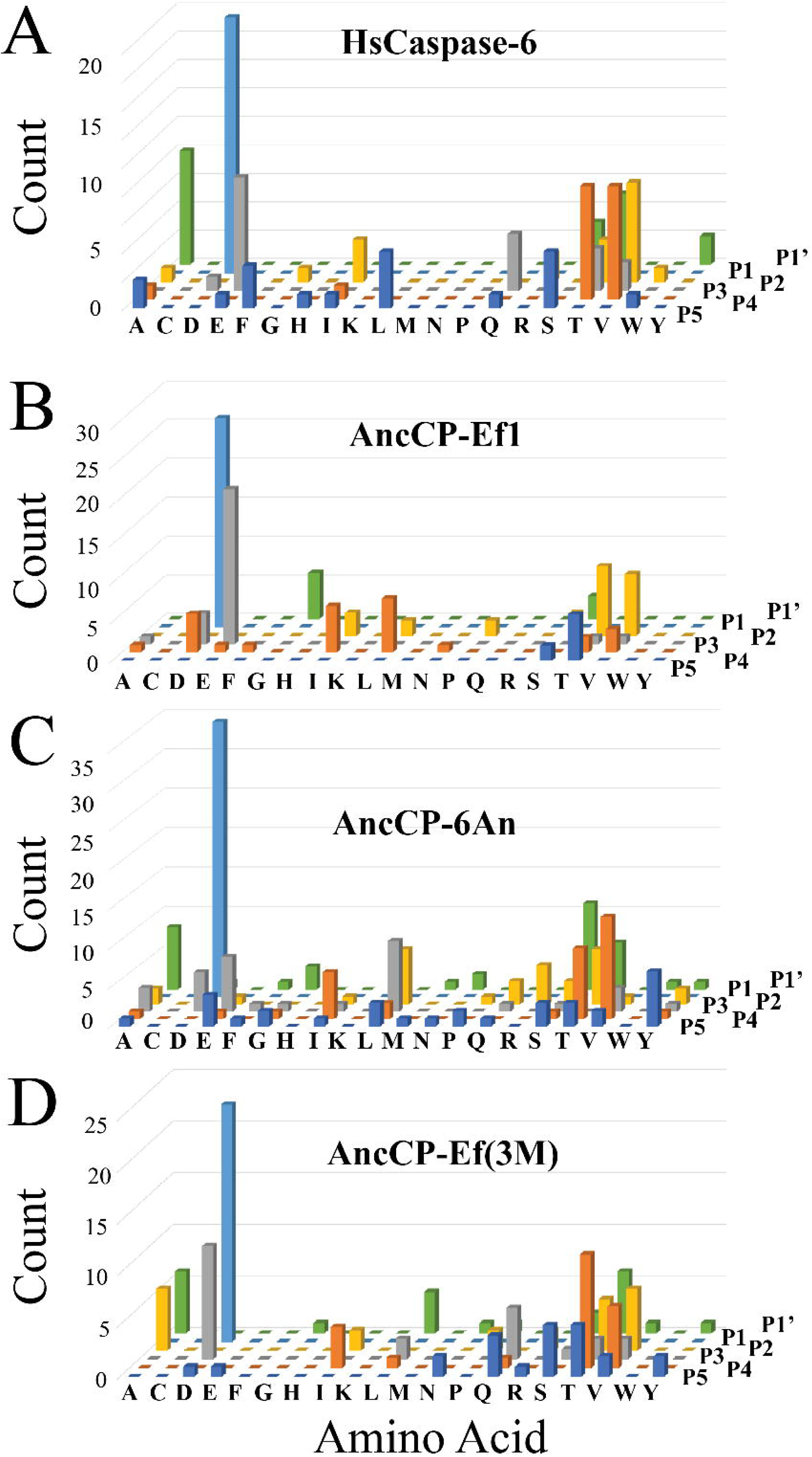
Substrate preferences determined by substrate-phage display. Amino acid preferences shown for substrate positions P5-P1’ for HsCaspase-6 (A), AncCP-Ef1 (B), AncCP-6An (C), and a triple mutant of AncCP-Ef1 (CP-N162E,CP-S172D,GP9-E01V). Values on the Y-axes indicate number of phage sequences containing the specified amino acid.

### Structures of ancestral caspases show evolutionary changes in enzyme selection

We determined the structures of AncCP-Ef1 and of AncCP-6An with either DEVD or VEID, respectively, bound in the active site. The proteins crystalized in the P2_1_2_1_2_1_ space group between 1.83Å and 2.61Å resolution (Supplemental Information, Table S4). The data show that the structures are very similar to extant caspases, with <0.5Å RMSD (Figure 4A), demonstrating that the caspase-hemoglobinase fold has been conserved for more than 650 million years. For AncCP-Ef1, active site loop 4 is partially disordered. For example, we observe no electron density for residues GP9-S03 to GP9-A08, although there is good electron density for the P4 Asp (Figure 4B). In the case of AncCP-6An (VEID), loop 4 and the P4 Val are well-ordered (Figs. 4C).

**Figure 4.**
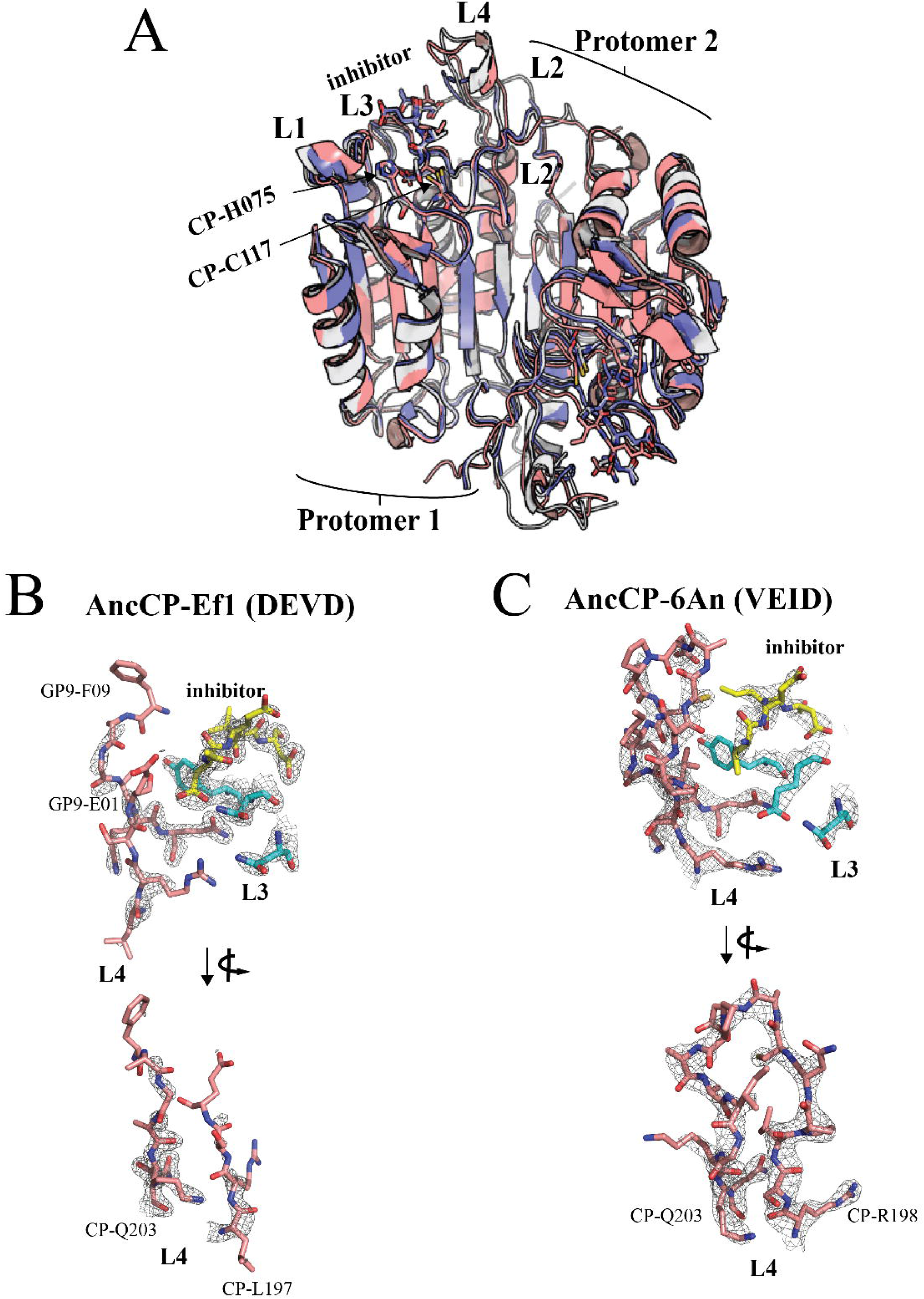
Structures of AncCP-Ef1 and of AncCP-6An. (A) Comparison of AncCP-Ef1 structure determined by X-ray crystallography (blue) to those of HsCaspase-3 (grey, PDB ID: 2J30) and AncCP-6An (peach). Total RMSD between the three structures is <0.5 Å. The five active site loops L1-L4 and L2’ are shown for protomer 1, and catalytic residues (CP-H075 and CP-C117) are indicated. (B-C) Electron density map (mesh) for active site loop 4 (L4, peach) and select amino acids of loop 3 (L3, blue) of AncCP-Ef1 (DEVD) (B) or AncCP-6An (VEID) (C) demonstrating regions of disorder in L4 of AnCP-Ef1 (panel B) compared to that of AncCP-6An (panel C), which is well-ordered.

In HsCaspase-3, the P4 Asp makes three hydrogen bonds with the side-chain of CP-N162, on active site loop L3, and to water molecules. The waters link the backbone atoms of GP9-E01 and of GP9-S02 to the P4 Asp (Figure 5A). As described above, CP-162 exhibits A1-type ambiguity since it is conserved as Asn in group II caspases or Glu in group III caspases (Figs. 5A,B). In order for Val to bind in the S4 pocket of HsCaspase-6, a network of four charged residues interact with CP-E162 so that it is positioned away from the P4 Val (CP-N165, CP-Q171, CP-D172, and CP-R198) (Figure 5B). Only one of the four charged residues are found in HsCaspase-3 (CP-Q171), although CP-165 is a conservative substitution (N to D).

**Figure 5.**
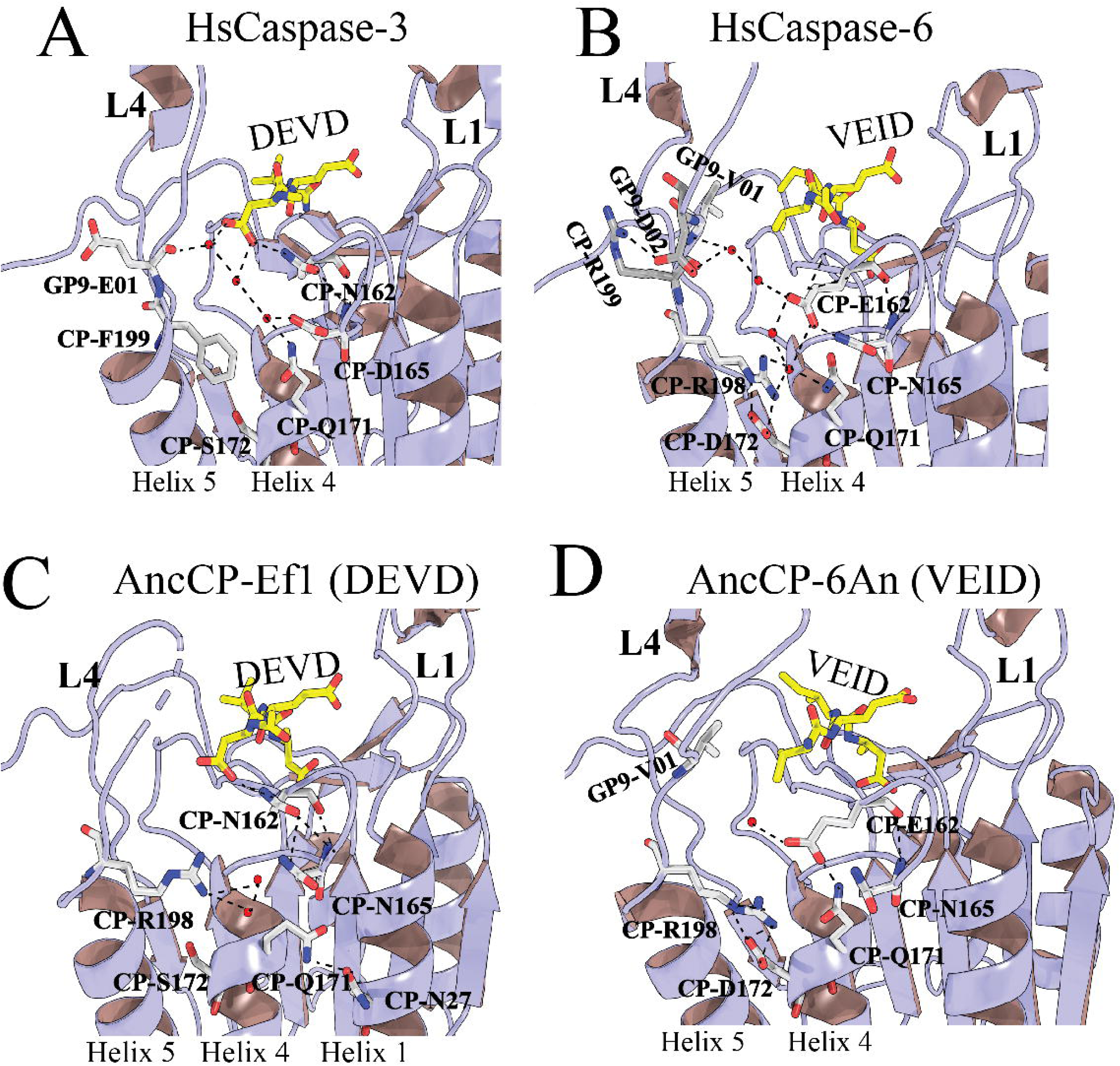
Electrostatic network near S4 subsite. Comparison of charged residue interactions near S4 for (A) HsCaspase-3 (PDB ID: 2J30) and (B) HsCaspase-6 (PDB ID: 3OD5). CP-N162 is conserved in caspase-3 and hydrogen bonds to the carboxylate of the substrate. In contrast, CP-E162 is conserved in caspase-6 and hydrogen bonds to CP-N165. Several polar residues (CP-Q171, CP-D172, CP-R198) form through-water hydrogen bonds with CP-E162. Three of the five polar residues (CPN165, CP-Q171, CP-R198) are present in AncCP-Ef1 (C). The five polar residues in HsCaspase-6 are observed in the common ancestor to caspase-6, AncCP-6An (D).

In AncCP-Ef1, CP-N162 interacts directly with the P4 aspartate, as in HsCaspase-3, but the hydrogen-bonding pattern is incomplete. We observe no hydrogen bonds with backbone atoms in loop 4, and CP-Q171 has rotated away from the active site and interacts with CP-N27 on Helix 1 (Figure 5C). In addition, CP-R198 is also rotated toward the P4 Asp. Thus, while AncCP-Ef1 contains three of the five amino acids in the network of HsCaspase-6, the hydrogen bonding network is not established. In contrast, the charged-network is fully formed in AncCP-6An, with the CP-E162 and CP-D172 substitutions, and the same H-bonding pattern is observed in AncCP-6An as in HsCaspase-6 (Figure 5D). Finally, we note that in HsCaspase-3, GP9-E01, on loop 4, is rotated away from the active site and toward solvent (Figure 5A). In caspase-6, the Glu is substituted with Val, which rotates toward the active site and forms part of the S4 binding pocket. In addition, CP-R199 and GP9-D09 flank the hydrophobic Val and form a salt bridge in HsCaspase-6 that may stabilize loop 4 (Figure 5B). In AncCP-Ef1 and AncCP-Ef2, one observes Glu at GP9-01, and the position changes to Val in AncCP-6An (Figure 1C). The CP-R199:GP9-D09 salt bridge is not observed until subsequent evolutionary nodes.

In order to examine the putative role of the charged network in substrate selection, we introduced several substitutions into AncCP-Ef1 and determined changes in enzyme activity on DEVD, VEID, and LETD tetrapeptide substrates. We first introduced substitutions at CP-162 (Asn to Glu) and GP9-01 (Glu to Val), as well as the double mutant (CP-N162E,GP9-E01V). The data show that activity increased ∼4-fold in the CP-N162E single mutant, but there was little effect on the selectivity (Supplemental Information, Table S5). In contrast, we observed a large decrease in activity (>10-fold) for the GP9-E01V single mutant. In this case, one observes changes in k_cat_ and in K_M_, resulting in the low activity. When the two substitutions were combined, however, the activity (k_cat_/K_M_) of the double mutant (CP-N162E,GP9-E01V) increased ∼10-fold for P4 Val or Leu and only ∼3-fold for P4 Asp. The increased selection was due to both an increase in k_cat_ and a decrease in K_M_ for the two hydrophobic P4 residues compared to P4 Asp (Figure 6). We observed the largest change in activity when we substituted CP-S172 with Asp along with the CP-N162E,GP9-E01V substitutions. The data for the triple mutant show ∼80-fold increase in activity for Val and ∼20-fold increase for Leu, with only ∼4-fold change for Asp at P4 (Figure 6A, Supplemental Information, Table S5). The increase in activity is largely due to a decrease in K_M_ (Figs. 6B,6C), which is below 10 μM for P4 Val. Overall, the series of mutants of AncCP-Ef1 show that the substitutions had little effect on the activity with P4 Asp (with the exception of the GP9-E01V single mutant). However, the three substitutions resulted in selection of Val over Asp through a large decrease in K_M_. In support of the tetrapeptide substrate data, we observed a similar shift in specificity using the substrate-phage display assay. In this case, the triple mutant of AncCP-Ef1 demonstrated a specificity (P5-P1’) of (S/T)(V/T)DVDA, with little or no selection for Leu or Asp at P4 (Figure 3D). We saw no further improvement in selection by including the CP-R199:GP9-D02 salt bridge with the three substitutions (Figure 6, Supplemental Information, Table S5), so the importance, if any, of the salt bridge in stabilizing loop 4 of caspase-6 is not yet clear. Overall, the data show that only three substitutions are required in AncCP-Ef1 to change substrate selection, where two substitutions complete the network of charged amino acids that position CP-E162 away from the S4 pocket while one substitution positions the hydrophobic GP9-V01 in the S4 pocket.

**Figure 6.**
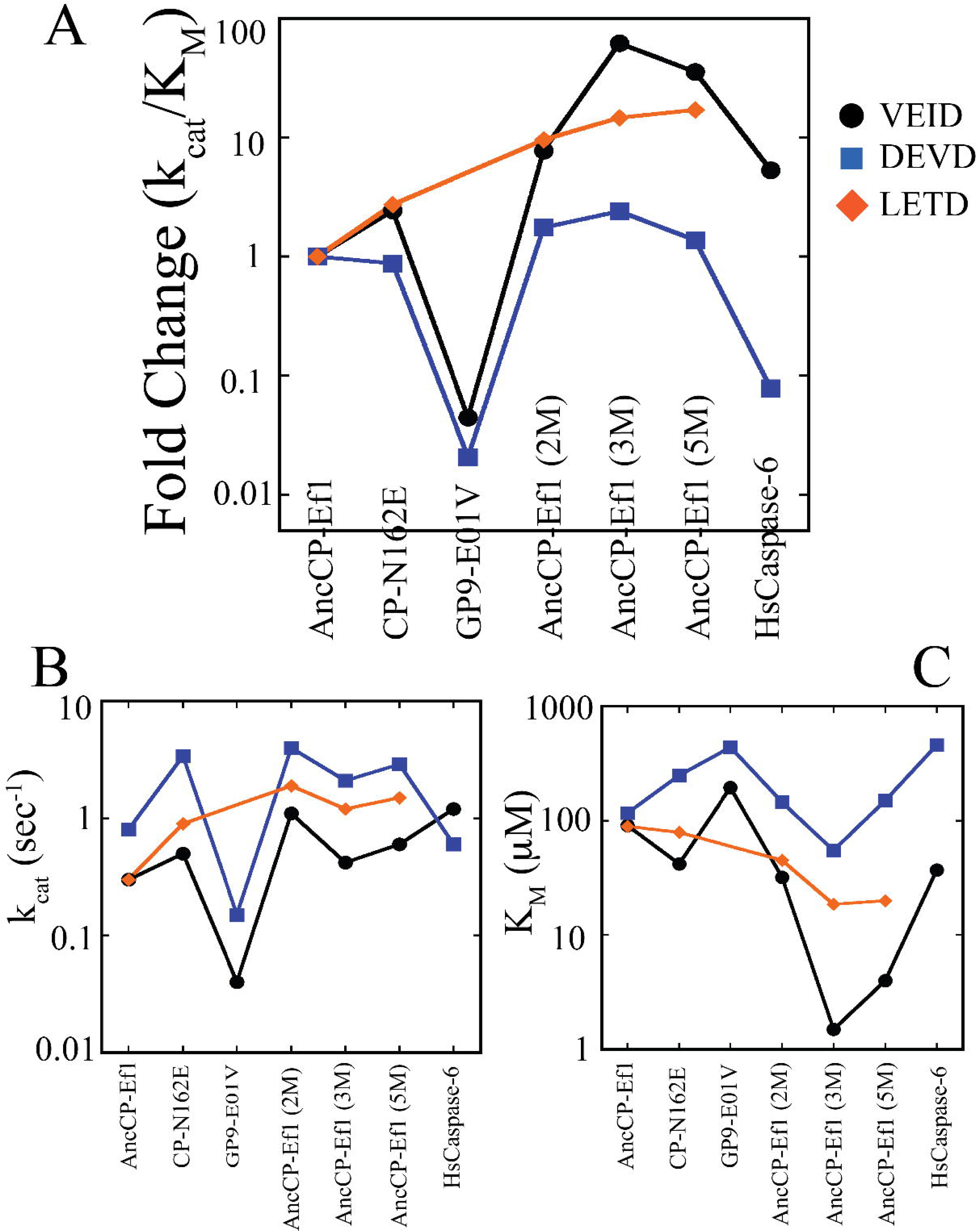
Enzyme activity of ancestral effector caspases. (A) Change in k_cat_/K_M_ for AncCP-Ef1 and several mutants for three substrates, VEID (black), DEVD (blue), and LETD (orange). Activity is normalized to that of AncCP-Ef1 in order to show relative increase in activity. Actual values of k_cat_ (B) and of K_M_ (C) are also shown for each protein. Single mutants (CP-N162E or GP9-E01V) refer to mutations in AncCP-Ef1. AncCP-Ef1 (2M), AncCP-Ef1 (3M), and AncCP-Ef1 (5M) refer to CP-N162E,GP9-E01V (2M), CP-N162E,CP-S172D,GP9-E01V (3M), and CP-N162E,CP-S172D,GP9-E01V, CP-S199R, GP9-S02D (5M). All enzymatic parameters are shown in Supplemental Information, Table S5.

### Variations in the C-terminus of helix-5

In caspase-6, the amino acid at CP-198 is a highly conserved arginine residue at the C-terminus of helix-5 at the junction with loop L4. As described above, the arginine participates in an interaction network that coordinates the charged carboxyl group of CP-E162 on loop L3 to position it away from the S4 pocket (Figure 5B), while also increasing the hydrophobicity by exposing the hydrophobic β and γ carbons of the side-chain of CP-E162 to the S4 pocket. The arginine at CP-198 (CP-R198) is also present in AncCP-Ef1; however, the interaction network in AncCP-Ef1 is missing a highly conserved aspartate residue on helix 4 observed in caspase-6 at CP-172, which forms hydrogen bonds with the side-chain of CP-R198. In contrast, the serine at CP-172 of AncCP-Ef1 is highly conserved in caspase-3, and the shorter side-chain does not interact with CP-E162. The substitutions in helix 5 result in structural changes in the last turn of the helix that affect loop L4 (Figure 7A). In HsCaspase-3 and in AncCP-Ef1, the backbone atoms of loop L4 are closer to the S4 binding pocket, and GP9-E01 is rotated toward solvent and away from the S4 pocket. In contrast, in HsCaspase-6 and AncCP-6An, the orientation of loop L4 is shifted by one residue as a result of the structural changes in helix 5 such that the backbone atoms are further from the S4 pocket. The shift causes GP9-V01 in HsCaspase-6 and in AncCP-6An to move into the S4 pocket.

**Figure 7.**
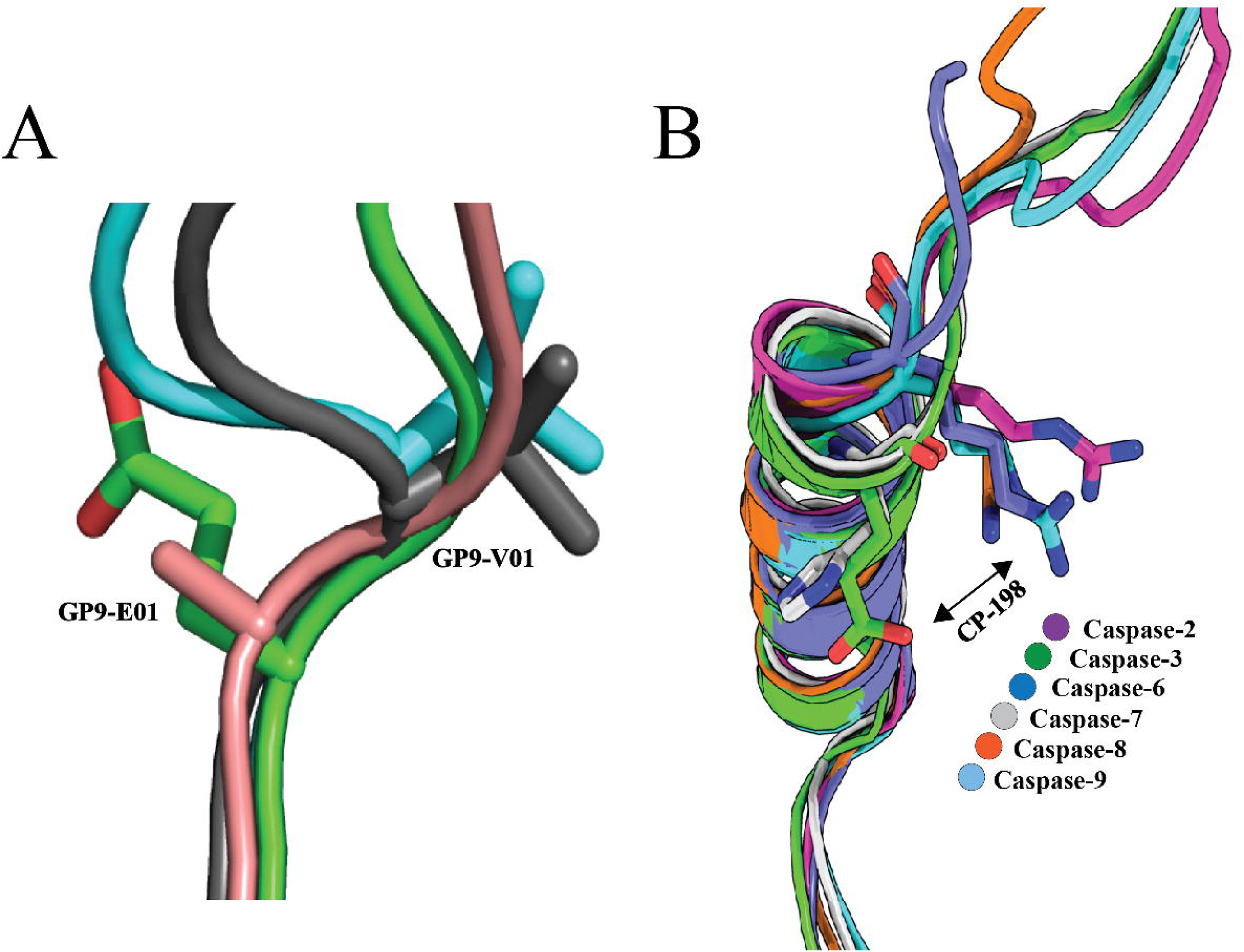
Changes in helix 5 and the adjacent active site loop 4. (A) Electrostatic network near S4 pocket in caspase-6 (cyan) results in repositioning GP9-V01 of loop 4 toward the S4 pocket. A similar orientation is observed for AncCP-6An (grey). In caspase-3 (green), the GP9-E01 side-chain is rotated toward solvent, and a similar orientation is observed in AncCP-Ef (peach). (B) Comparison of helix 5 and loop 4 of extant enzymes show a change in the final turn of helix 5 for all caspases except caspases-3 and -7, which repositions CP-198 toward the S4 binding pocket.

An analysis of initiator caspases revealed that caspase-2 also has a highly conserved arginine at CP-198, while caspases-8 and -9 utilize lysine residues in the same position. The structures of the initiator caspases show the same orientation of the helix 5-loop L4 residues as observed for HsCaspase-6 and AncCP-6An (Figure 7B). Comparatively, caspases-3 and -7 are outliers. The CP-E198 (caspase-3) or CP-H198 (caspase-7), as well as the more regular helical turn, orient the charged groups away from loop L3 and the charged interaction network observed in HsCaspase-6. Therefore, caspases that recognize hydrophobic residues (I/L/W/V) in the P4 substrate position utilize the long positively charged side-chain at CP-198 and subsequent alterations to the final turn of helix 5. We note that the interaction of CP-R198 and CP-D172 is uniformly conserved in all chordate caspase-6 genes.

## Discussion

Since the pioneering work of Zuckerkandl and Pauling over 50 years ago [67, 68], research in the evolutionary biochemistry of protein structure-function relationships has aimed to apply rigorous biochemical and biophysical methods to understand how protein sequence changes affect protein structure and function. Understanding how random chance, selection pressure, and changes in free-energy landscapes determine the characteristics of divergent proteins within a family allows one to track form and function along a phylogeny [41]. Specific, and limited, combinations of amino acids gleaned from comparative studies of extant proteins do not include the historical context that may allow the combination of amino acids to function in the active site environment. Ancestral protein reconstruction (APR) adds the dimension of evolutionary time and fitness to the structure-function relationship [63, 64] and provides powerful tools to characterize evolutionary changes in proteins experimentally [69–71]. The evolutionary trajectory is thus determined by a combination of biochemical, biophysical, and regulatory factors, but the interplay among these factors and their role in protein evolution remains an unresolved question.

We used APR techniques to infer ancestral sequences and to resurrect the common ancestor of effector caspases-3/6/7 and of caspase-6. The analysis provides site-specific probabilities throughout the protein sequence such that one can examine the robustness of the APR methods by characterizing multiple proteins from the pool of possible sequences in an ancestral node [64]. Ambiguity, or uncertainty in ancestral sequence inference, is typically attributed to insufficient sequence data, uncertain gap placement in the multiple sequence alignment, tree topology, and the extent of sequence divergence relative to tree articulation [64, 72]. As we described previously [42], the large dataset compilation from the CaspBase minimizes the major causes of ambiguity. Regions with higher ambiguity are generally observed in sites that are not evolutionarily constrained for proper structure-function, such as surface exposed residues, loops, turns, and intrinsically disordered regions [73]. In addition, assessing data from multiple reconstructions provides insight about positions of interest. For example, we resurrected two proteins from the common ancestral node (AncCP-Ef1 and AncCP-Ef2) and showed similar characteristics in that both enzymes had low activity and little selection against P4 valine, aspartate, or leucine, suggesting that both proteins robustly represent the pool of possible proteins in the ancestral node. Amino acid positions that were ambiguous were mostly found in the surface helices as well as two regions that are flexible or disordered, the intersubunit linker and prodomain (type A2 ambiguity). The surface helices were previously characterized as sites of allosteric regulation [22,60,61], so our data suggest that allosteric sites may have different evolutionary pressures compared to the active site residues. That is, the allosteric sites may have evolved in a species-dependent manner based on individual needs, leading to larger sequence variations at those sites and higher ambiguity in the APR analysis. Overall, our data agree with previous studies showing that the pool of possible ancestors at the nodes of a phylogenetic tree reflect ancestral mechanistic function experimentally [74–76].

The structure of AncCP-Ef1 with DEVD bound in the active site showed that the caspase-hemoglobinase fold has been conserved for over 650 million years as the protein exhibits <0.5 Å RMSD with extant HsCaspase-3. The data also showed that active site loop L4 was partially disordered in the ancestral effector caspase. In contrast, loop 4 was well-formed in the ancestral caspase-6 enzyme, AncCP-6An, with VEID inhibitor bound in the active site. For AncCP-Ef1 (DEVD), the hydrogen-bonding network with the P4 Asp, observed in HsCaspase-3, was incomplete in that interactions were missing between loop L4 and the P4 Asp while interactions with loop L3 were retained. The multiple structures showed flexibility in the side chains of CP-R198 and CP-Q171 as the side-chains were observed in different orientations. In AncCP-6An, both amino acids are part of a network of charged residues between loop L3, helix 5, and loop L4 that ultimately interact with CP-E162 to position the charged side-chain away from the hydrophobic S4 pocket. In AncCP-Ef1 and HsCaspase-3, the incomplete network, as well as the shorter side-chain of CP-N162, positions CP-N162 to interact directly with the P4 Asp and thus retains the hydrophilic character of the S4 pocket. Two of the five residues in the network are not found in AncCP-Ef1, so we completed the network by substituting the two sites (CP-S172D, CP-N162E), as well as introducing GP9-V01. In HsCaspase-6 and in AncCP-6An, the substitution of GP9-E01V results in rotation of the valine into the S4 pocket. Overall, the data for the triple mutant showed ∼80-fold increase in the activity against VEID substrate with little change in the activity for DEVD. The activity increase was due primarily to a large decrease in K_M_, suggesting that the substitutions resulted in improved binding of P4 Val without affecting the binding of P4 Asp. Thus, with only three substitutions in the ancestral effector caspase scaffold, the enzyme became selective for P4 Val *versus* Asp.

Extant genes accumulate mutations as described by neutral mutation theory [77], and their probability of fixation into a population is described by models of genetic drift [78]. A few outcomes can occur for nascent genes, but the majority of mutations are deleterious and will likely cause loss of function, or pseudogenization, which leads to purification from the population [79]. In the ancestral effector caspase and ancestral caspase-6, only three vertical substitutions of the forty-two evolutionary changes resulted in altered substrate specificity in the caspase-6 lineage. The remaining vertical mutations may contribute to protein epistasis or to the evolution of allosteric sites. Together the data show that enzyme specificity was established early in the evolution of caspase-6 and that allosteric regulation likely followed through subsequent evolution. Intriguingly, the data suggest that features of the conformational landscape in the ancestral effector caspase remain in extant enzymes. Because so few substitutions to the conserved scaffold were required for neofunctionalization of substrate specificity (three substitutions of ∼260 amino acids), subsequent mutations may then affect access to various conformational states through changes in the free-energy landscape. If this is true, then the unique coil-to-helix transition observed in caspase-6, for example, may still be present in the conformational landscapes of other caspases but is inaccessible due to evolutionary changes that introduced high barriers to the state.

Understanding the evolutionary changes in conformational landscapes that resulted in substrate selection and in allosteric regulation may provide strategies for re-engineering caspases with desired substrate selection coupled to unique conformational states. Our APR studies show that the methodology is effective for characterizing evolutionary changes in ancestral proteins in order to infer functional changes in the extant caspase proteases. By focusing on peptide mutations (substitutions, insertions, and deletions) to quantify the probability of evolutionary change, the APR methodology allows for targeted assays by examining the fewer positions that changed between evolutionary nodes, as compared to the larger substitutions between extant enzymes.

## Author contributions

RDG, SS, JT-M, PDS, and ACC designed the experiments; RDG, SS, and JT-M carried out the experiments. RDB and ACC wrote the manuscript, and all authors contributed to data analysis. All authors have approved the manuscript.

## Funding

This work was supported by a grant from the National Institutes of Health [grant number GM127654 (to A.C.C)] and by funds from UT Arlington [Office of the Vice President for Research (to A.C.C)]. Use of the Advanced Photon Source was supported by the U.S. Department of Energy, Office of Science, Office of Basic Energy Sciences, under contract number W-31-109-ENG-38.

## Competing interests

The authors declare that there are no competing interests associated with this manuscript

## References

1 Bayles, K. W. (2014) Bacterial programmed cell death: Making sense of a paradox. Nat. Rev. Microbiol. 12, 63–69.

2 Zmasek, C. M., Zhang, Q., Ye, Y. and Godzik, A. (2007) Surprising complexity of the ancestral apoptosis network. Genome Biol. 8, R226.

3 Crawford, E. D., Seaman, J. E., Barber, A. E., David, D. C., Babbitt, P. C., Burlingame, A. L. and Wells, J. A. (2012) Conservation of caspase substrates across metazoans suggests hierarchical importance of signaling pathways over specific targets and cleavage site motifs in apoptosis. Cell Death Differ. 19, 2040– 2048.

4 Allocati, N., Masulli, M., Di Ilio, C. and De Laurenzi, V. (2015) Die for the community: An overview of programmed cell death in bacteria. Cell Death Dis. 6, e1609.

5 Cortesi, F., Musilová, Z., Stieb, S. M., Hart, N. S., Siebeck, U. E., Malmstrøm, M., Tørresen, O. K., Jentoft, S., Cheney, K. L., Marshall, N. J., et al. (2015) Ancestral duplications and highly dynamic opsin gene evolution in percomorph fishes. Proc. Natl. Acad. Sci. 112, 1493–1498.

6 Eckhart, L., Ballaun, C., Hermann, M., VandeBerg, J. L., Sipos, W., Uthman, A., Fischer, H. and Tschachler, E. (2008) Identification of novel mammalian caspases reveals an important role of gene loss in shaping the human caspase repertoire. Mol Biol Evol 25, 831–841.

7 Boatright, K. M. and Salvesen, G. S. (2003) Mechanisms of caspase activation. Curr. Opin. Cell Biol. 15, 725–731.

8 Thomsen, N. D., Koerber, J. T. and Wells, J. A. (2013) Structural snapshots reveal distinct mechanisms of procaspase-3 and -7 activation. Proc. Natl. Acad. Sci. 110, 8477–8482.

9 Clark, A. C. (2016) Caspase allostery and conformational selection. Chem. Rev 116, 6666–6706.

10 Oberst, A., Dillon, C. P., Weinlich, R., McCormick, L. L., Fitzgerald, P., Pop, C., Hakem, R., Salvesen, G. S. and Green, D. R. (2011) Catalytic activity of the caspase-8-FLIP(L) complex inhibits RIPK3-dependent necrosis. Nature 471, 363– 367.

11 Muzio, M., Stockwell, B. R., Stennicke, H. R., Salvesen, G. S. and Dixit, V. M. (1998) An induced proximity model for caspase-8 activation. J. Biol. Chem. 273, 2926–2930.

12 Dagbay, K., Eron, S. J., Serrano, B. P., Velázquez-Delgado, E. M., Zhao, Y., Lin, D., Vaidya, S. and Hardy, J. a. (2014) A multipronged approach for compiling a global map of allosteric regulation in the apoptotic caspases. Methods Enzymol. 544, 215–49.

13 Penning, T. M. and Jez, J. M. (2001) Enzyme redesign. Chem. Rev. 101, 3027– 3046.

14 Cera, E. D. I. and Cantwell, A. M. (2001) Determinants of thrombin specificity. Ann. New York Acad. Sci. 936, 133–146.

15 MacGregor, E. A., Janecek, S. and Svensson, B. (2001) Relationship of sequence and structure to specificity in the a-amylase family of enzymes. Biochem. Biophys. Acta 1546, 1–20.

16 Mildvan, A. S. (1974) Mechanism of enzyme action. Annu. Rev. Biochem. 43, 357–399.

17 Bender, M. L. and Kezdy, F. J. (1965) Mechanism of action of proteolytic enzymes. Annu. Rev. Biochem. 34, 49–76.

18 Mitchell, J. B. (2017) Enzyme function and its evolution. Curr. Opin. Struct. Biol. 47, 151–156.

19 Pabis, A., Risso, V. A., Sanchez-Ruiz, J. M. and Kamerlin, S. C. (2018) Cooperativity and flexibility in enzyme evolution. Curr. Opin. Struct. Biol. 48, 83– 92.

20 Poreba, M., Strózyk, A., Salvesen, G. S. and Drag, M. (2013) Caspase substrates and inhibitors. Cold Spring Harb. Perspect. Biol. 5, 1–20.

21 Denault, J.-B. and Salvesen, G. S. (2003) Human caspase-7 activity and regulation by its N-terminal peptide. J. Biol. Chem. 278, 34042–34050.

22 Thomas, M. E., Grinshpon, R., Swartz, P. and Clark, A. C. (2018) Modifications to a common phosphorylation network provide individualized control in caspases. J. Biol. Chem. 293, 5447–5461.

23 MacPherson, D. J., Mills, C. L., Ondrechen, M. J. and Hardy, J. A. (2019) Tri-arginine exosite patch of caspase-6 recruits substrates for hydrolysis. J. Biol. Chem. 294, 71–88.

24 Hardy, J. A., Lam, J., Nguyen, J. T., O’Brien, T. and Wells, J. A. (2004) Discovery of an allosteric site in the caspases. Proc. Natl. Acad. Sci. USA 101, 12461–12466.

25 Scheer, J. M., Romanowski, M. J. and Wells, J. A. (2006) A common allosteric site and mechanism in caspases. Proc. Natl. Acad. Sci. USA 103, 7595–7600.

26 Talanian, R. V, Quinlan, C., Trautz, S., Hackett, M. C., Mankovich, J. A., Banach, D., Ghayur, T., Brady, K. D. and Wong, W. W. (1997) Substrate specificites of caspase family proteases. J. Biol. Chem. 272, 9677–9682.

27 Wejda, M., Impens, F., Takahashi, N., Van Damme, P., Gevaert, K. and Vandenabeele, P. (2012) Degradomics reveals that cleavage specificity profiles of caspase-2 and effector caspases are alike. J. Biol. Chem. 287, 33983–33995.

28 Ah, Y. L., Byoung, C. P., Jang, M., Cho, S., Do, H. L., Sang, C. L., Pyung, K. M. and Sung, G. P. (2004) Identification of caspase-3 degradome by two-dimensional gel electrophoresis and matrix-assisted laser desorption/ionization-time of flight analysis. Proteomics 4, 3429–3436.

29 Lüthi, A. U. and Martin, S. J. (2007) The CASBAH: a searchable database of caspase substrates. Cell Death Differ 14, 641–650.

30 Hill, M. E., Macpherson, D. J., Wu, P., Julien, O., Wells, J. A. and Hardy, J. A. (2016) Reprogramming caspase-7 specificity by regio-specific mutations and selection provides alternate solutions for substrate recognition. Chem. Biol. 11, 1603–1612.

31 Gulyaeva, N. V. (2003) Non-apoptotic functions of caspase-3 in nervous tissue. Biochem. 68, 1171–1180.

32 Iijima, N. and Yokoyama, T. (2007) Apoptosis in the medaka embryo in the early developmental stage. Acta Histochem. Cytochem. 40, 1–7.

33 Grotzer, M. A., Eggert, A., Zuzak, T. J., Janss, A. J., Marwaha, S., Wiewrodt, B. R., Ikegaki, N., Brodeur, G. M. and Phillips, P. C. (2000) Resistance to TRAIL-induced apoptosis in primitive neuroectodermal brain tumor cells correlates with a loss of caspase-8 expression. Oncogene 19, 4604–4610.

34 Hyman, B. T. and Yuan, J. (2012) Apoptotic and non-apoptotic roles of caspases in neuronal physiology and pathophysiology. Nat Rev Neurosci 13, 395–406.

35 Jacob, F. (1977) Evolution and tinkering. Science 196, 1161–1166.

36 Jensen, R. A. (1976) Enzyme recruitment in evolution of new function. Annu. Rev. Microbiol. 30, 409–425.

37 Khersonsky, O. and Tawfik, D. S. (2010) Enzyme promiscuity: a mechanistic and evolutionary perspective. Annu. Rev. Biochem. 79, 471–505.

38 Gerlt, J. A. and Babbitt, P. C. (2009) Enzyme (re)design: lessons from natural evolution and computation. Curr. Opin. Chem. Biol. 13, 10–18.

39 Sikosek, T. and Chan, H. S. (2014) Biophysics of protein evolution and evolutionary protein biophysics. J. R. Soc. Interface 11, 20140419–20140419.

40 Figliuzzi, M., Jacquier, H., Schug, A., Tenaillon, O. and Weigt, M. (2016) Coevolutionary landscape inference and the context-dependence of mutations in beta-lactamase tem-1. Mol. Biol. Evol. 33, 268–280.

41 Harms, M. J. and Thornton, J. W. (2013) Evolutionary biochemistry: Revealing the historical and physical causes of protein properties. Nat. Rev. Genet. 14, 559– 571.

42 Grinshpon, R. D., Williford, A., Titus-McQuillan, J. and Clark, A. C. (2018) The CaspBase: a curated database for evolutionary biochemical studies of caspase functional divergence and ancestral sequence inference. Protein Sci. 27, 1857– 1870.

43 Pei, J. and Grishin, N. V. (2007) PROMALS: towards accurate multiple sequence alignments of distantly related proteins. Bioinformatics 23, 802–808.

44 Pei, J. and Grishin, N. V. (2014) PROMALS3D: multiple protein sequence alignment enhanced with evolutionary and 3-dimensional structural information. Methods Mol. Biol. 1079, 263–271.

45 Kearse, M., Moir, R., Wilson, A., Stones-Havas, S., Cheung, M., Sturrock, S., Buxton, S., Cooper, A., Markowitz, S., Duran, C., et al. (2012) Geneious Basic: An integrated and extendable desktop software platform for the organization and analysis of sequence data. Bioinformatics 28, 1647–1649.

46 Darriba, D., Taboada, G. L., Doallo, R. and Posada, D. (2011) ProtTest-HPC: Fast selection of best-fit models of protein evolution. Bioinformatics 27, 1164–1165.

47 Burnham, K. P. and Anderson, D. R. (2002) Model Selection and Multimodel Inference: A practical Information-theoretic Approach (Burnham, K. P., and Anderson, D. R., eds.) 2nd ed., Springer-Verlag, New York.

48 Nguyen, L. T., Schmidt, H. A., Von Haeseler, A. and Minh, B. Q. (2015) IQ-TREE: A fast and effective stochastic algorithm for estimating maximum-likelihood phylogenies. Mol. Biol. Evol. 32, 268–274.

49 Felsenstein, J. (1985) Confidence limits on phylogenies: An approach using the bootstrap. Evolution 39, 783–791.

50 Ashkenazy, H., Penn, O., Doron-Faigenboim, A., Cohen, O., Cannarozzi, G., Zomer, O. and Pupko, T. (2012) FastML: A web server for probabilistic reconstruction of ancestral sequences. Nucleic Acids Res. 40, 580–584.

51 Le, S. Q. and Gascuel, O. (2008) An improved general amino acid replacement matrix. Mol. Biol. Evol. 25, 1307–1320.

52 Vaidya, S., Velázquez-Delgado, E. M., Abbruzzese, G. and Hardy, J. A. (2011) Substrate-induced conformational changes occur in all cleaved forms of caspase-6. J. Mol. Biol. 406, 75–91.

53 Feeney, B. and Clark, A. C. (2005) Reassembly of active caspase-3 is facilitated by the propeptide. J. Biol. Chem. 280, 39772–39785.

54 Bose, K., Pop, C., Feeney, B. and Clark, A. C. (2003) An uncleavable procaspase-3 mutant has a lower catalytic efficiency but an active site similar to that of mature caspase-3. Biochemistry 42, 12298–12310.

55 Pop, C., Feeney, B., Tripathy, A. and Clark, A. C. (2003) Mutations in the procaspase-3 dimer interface affect the activity of the zymogen. Biochemistry 42, 12311–12320.

56 Gasteiger, E., Hoogland, C., Gattiker, A., Duvaud, S., Wilkins, M. R., Appel, R. D. and Bairoch, A. (2005) Protein Analysis Tools on the ExPASy Server. In The proteomics protocols handbook: Protein Identification and Analysis Tools on the ExPASy server (Walker, J. M., ed.), pp 571–607, Humana Press, Totowa.

57 Tucker, M. B., MacKenzie, S. H., Maciag, J. J., Dirscherl Ackerman, H., Swartz, P., Yoder, J. A., Hamilton, P. T. and Clark, A. C. (2016) Phage display and structural studies reveal plasticity in substrate specificity of caspase-3a from zebrafish. Protein Sci. 25, 2076–2088.

58 Feeney, B., Pop, C., Tripathy, A. and Clark, A. C. (2004) Ionic interactions near the loop L4 are important for maintaining the active-site environment and the dimer stability of (pro)caspase 3. Biochem. J. 384, 515–25.

59 Feeney, B., Pop, C., Swartz, P., Mattos, C. and Clark, A. C. (2006) Role of loop bundle hydrogen bonds in the maturation and activity of (pro)caspase-3. Biochemistry 45, 13249–13263.

60 Vaidya, S. and Hardy, J. A. (2011) Caspase-6 latent state stability relies on helical propensity. Biochemistry 50, 3282–3287.

61 Maciag, J. J., Mackenzie, S. H., Tucker, M. B., Schipper, J. L., Swartz, P. and Clark, A. C. (2016) Tunable allosteric library of caspase-3 identifies coupling between conserved water molecules and conformational selection. Proc. Natl. Acad. Sci. 113, 6080–6088.

62 Wang, X.-J., Cao, Q., Liu, X., Wang, K.-T., Mi, W., Zhang, Y., Li, L.-F., LeBlanc, A. C. and Su, X.-D. (2010) Crystal structures of human caspase 6 reveal a new mechanism for intramolecular cleavage self-activation. EMBO Rep. 11, 841–847.

63 Harms, M. J. and Thornton, J. W. (2010) Analyzing protein structure and function using ancestral gene reconstruction. Curr. Opin. Struct. Biol. 20, 360–366.

64 Eick, G. N., Bridgham, J. T., Anderson, D. P., Harms, M. J. and Thornton, J. W. (2017) Robustness of reconstructed ancestral protein functions to statistical uncertainty. Mol. Biol. Evol. 34, 247–261.

65 Roschitzki-Voser, H., Schroeder, T., Lenherr, E. D., Frölich, F., Schweizer, A., Donepudi, M., Ganesan, R., Mittl, P. R. E., Baici, A. and Grütter, M. G. (2012) Human caspases in vitro: Expression, purification and kinetic characterization. Protein Expr. Purif. 84, 236–246.

66 Tang, Y., Wells, J. A. and Arkin, M. R. (2011) Structural and enzymatic insights into caspase-2 protein substrate recognition and catalysis. J. Biol. Chem. 286, 34147–34154.

67 Zuckerkandl, E. and Pauling, L. (1965) Molecules as documents of evolutionary biology. J. Theor. Biol. 8, 357–366.

68 Zuckerkandl, E. and Pauling, L. (1965) Evolutionary divergence and convergence in proteins. In Evolving Genes and Proteins (Bryson, V., and Vogel, H. J., eds.), pp 97–166, Academic Press, New York.

69 Newton, M. S., Arcus, V. L., Gerth, M. L. and Patrick, W. M. (2018) Enzyme evolution: innovation is easy, optimization is complicated. Curr. Opin. Struct. Biol. 48, 110–116.

70 Johansson, K. E. and Lindorff-Larsen, K. (2018) Structural heterogeneity and dynamics in protein evolution and design. Curr. Opin. Struct. Biol. 48, 157–163.

71 Capra, E. J., Perchuk, B. S., Skerker, J. M. and Laub, M. T. (2012) Adaptive mutations that prevent crosstalk enable the expansion of paralogous signaling protein families. Cell 150, 222–232.

72 Duchêne, S. and Lanfear, R. (2015) Phylogenetic uncertainty can bias the number of evolutionary transitions estimated from ancestral state reconstruction methods. J. Exp. Zool. Part B Mol. Dev. Evol. 324, 517–524.

73 Hanson-Smith, V., Kolaczkowski, B. and Thornton, J. W. (2010) Robustness of ancestral sequence reconstruction to phylogenetic uncertainty. Mol. Biol. Evol. 27, 1988–1999.

74 Howard, C. J., Hanson-Smith, V., Kennedy, K. J., Miller, C. J., Lou, H. J., Johnson, A. D., Turk, B. E. and Holt, L. J. (2014) Ancestral resurrection reveals evolutionary mechanisms of kinase plasticity. Elife 3, 1–22.

75 Wheeler, L. C., Lim, S. A., Marqusee, S. and Harms, M. J. (2016) The thermostability and specificity of ancient proteins. Curr. Opin. Struct. Biol. 38, 37– 43.

76 Babkova, P., Sebestova, E., Brezovsky, J., Chaloupkova, R. and Damborsky, J. (2017) Ancestral haloalkane dehalogenases show robustness and unique substrate specificity. ChemBioChem 18, 1448–1456.

77 Kimura, M. (1968) Evolutionary rate at the molecular level. Nature 217, 624–626.

78 Lynch, M., Ackerman, M. S., Gout, J. F., Long, H., Sung, W., Thomas, W. K. and Foster, P. L. (2016) Genetic drift, selection and the evolution of the mutation rate. Nat. Rev. Genet. 17, 704–714.

79 Zhang, J. (2003) Evolution by gene duplication: An update. Trends Ecol. Evol. 18, 292–298.

