## Supplemental Information for "Resurrection of ancestral effector caspases identifies novel networks for evolution of substrate specificity"

#### Supplemental Fig. Legends

**Supplemental Figure. S1.** Amino acids in S1 (A), S2 (B), S3 (C), and S4 (D) binding pockets for caspase-6 (PDB ID: 3OD5). Amino acids in each site are shown by stick and semi-transparent sphere, inhibitor is shown in yellow, and P4-P1 residues of the inhibitor (VEID) are labeled in red.

**Supplemental Figure. S2.** Comparison of ancestral and human effector caspases percent amino acid identity.

**Supplemental Figure. S3.** Site-specific differences (orange) between AncCP-Ef1 and Anc CP-Ef2.

Supplemental Figure 1

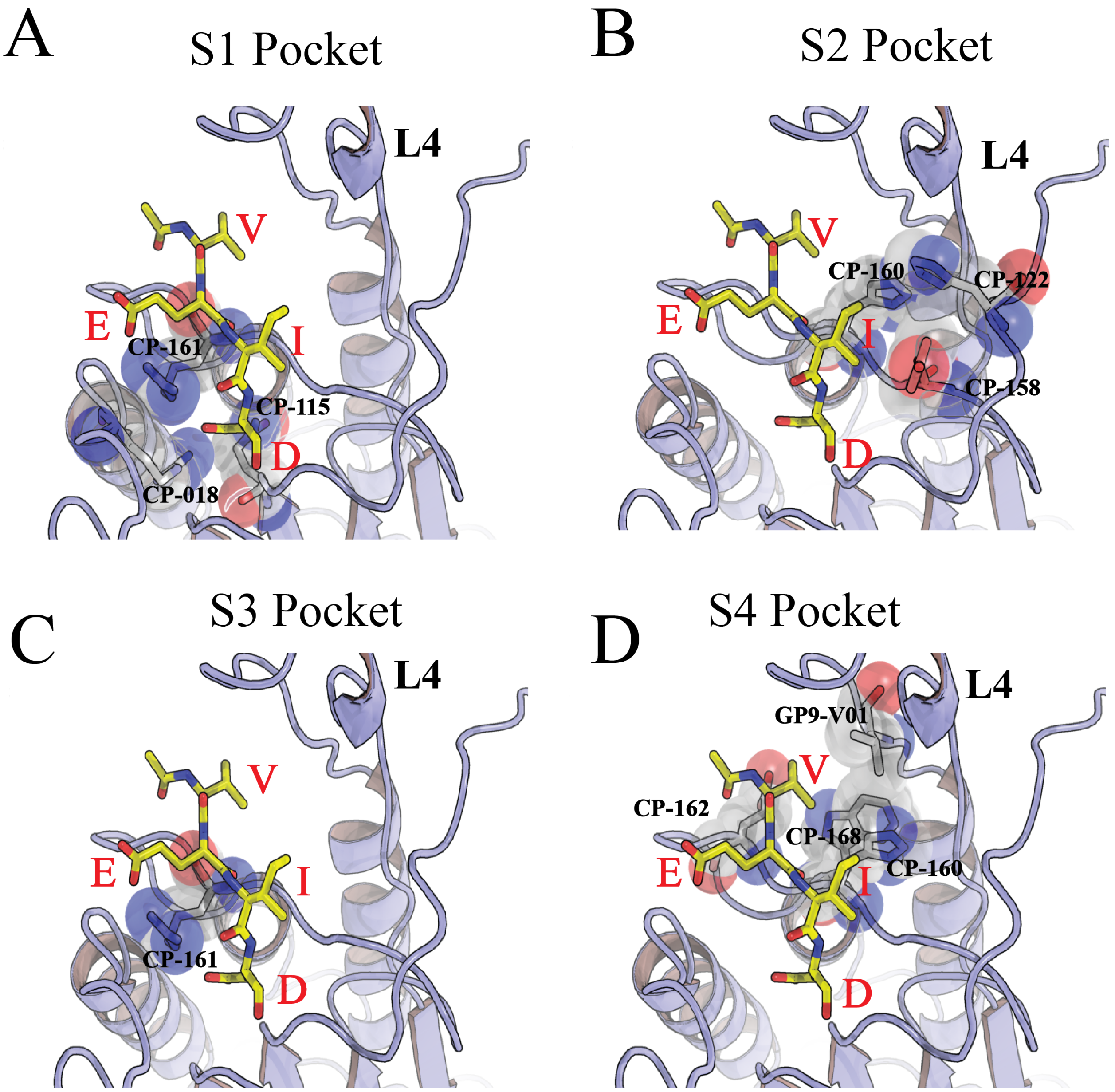

Supplemental Figure 2

| | AncCP-Ef1 | AncCP-6An | HsCaspase-6 | HsCaspase-3 | HsCaspase-7 |
|---|---|---|---|---|---|
| AncCP-Ef1 | | 68 | 54 | 60 | 58 |
| AncCP-6An | | | 75 | 45 | 43 |
| HsCaspase-6 | | | | 40 | 37 |
| HsCaspase-3 | | | | | 56 |
| HsCaspase-7 | | | | | |

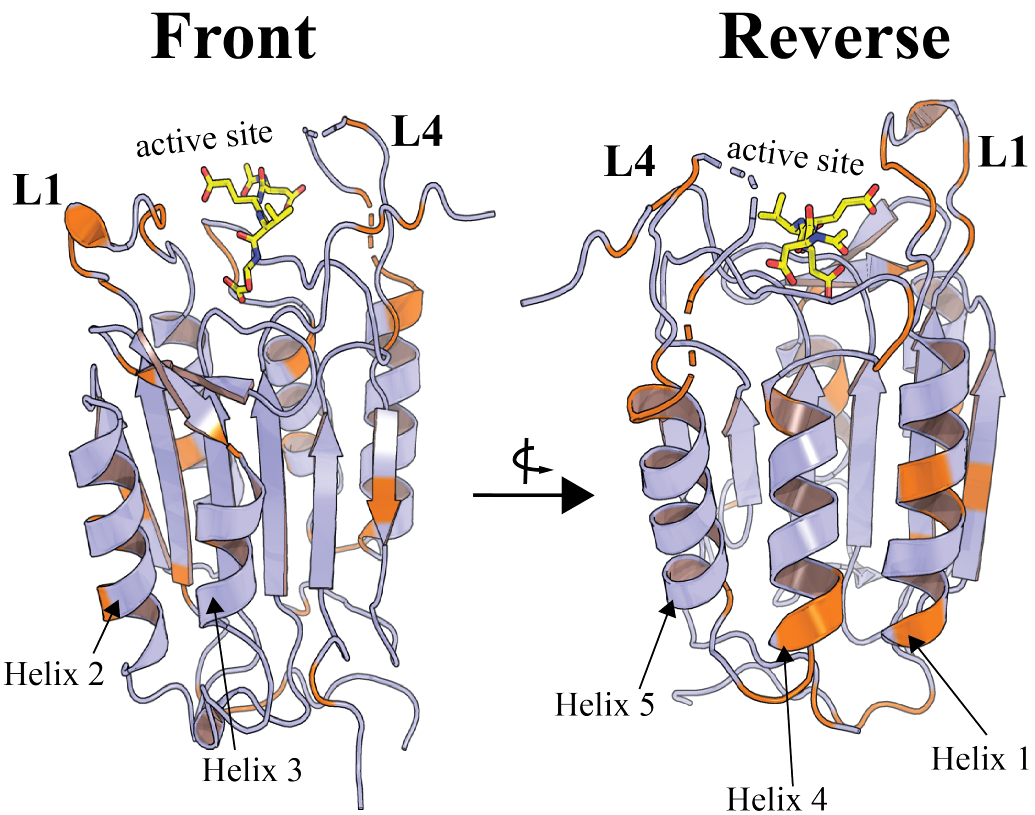
Front
Reverse
active site
L1
L4
Helix 2
Helix 3
active site
L4
L1
Helix 5
Helix 4
Helix 1

**Supplemental Table S1. APR_1: List of Taxa and caspases used in reconstructions from initial caspase data base (pre-CaspBase).**

| Caspase | Species |
|---|---|
| Caspase-1 | Anolis carolinensis |
| Caspase-1 | Camelus bactrianus |
| Caspase-1 | Danio rerio |
| Caspase-1 | Equus caballus |
| Caspase-1 | Gallus gallus |
| Caspase-1 | Melopsittacus undulatus |
| Caspase-10 | Acinonyx jubatus |
| Caspase-10 | Aptenodytes forsteri |
| Caspase-10 | Cariama cristata |
| Caspase-10 | Carlito syrichta |
| Caspase-10 | Chelonia mydas |
| Caspase-10 | Chrysochloris asiatica |
| Caspase-10 | Equus caballus |
| Caspase-10 | Erinaceus europaeus |
| Caspase-10 | Gallus gallus |
| Caspase-10 | Nestor notabilis |
| Caspase-10 | Pongo abelii |
| Caspase-10 | Taeniopygia guttata |
| Caspase-10 | Tinamus guttatus |
| Caspase-10 | Xenopus tropicalis |
| Caspase-10-isoform-X1 | Anolis carolinensis |
| Caspase-10-isoform-X1 | Balaenoptera acutorostrata scammoni |
| Caspase-10-isoform-X1 | Odobenus rosmarus divergens |
| Caspase-12-isoform-X1 | Pongo abelii |
| Caspase-13-like | Equus caballus |
| Caspase-13-like | Erinaceus europaeus |

| | |
|---|---|
| Caspase-13-like | Latimeria chalumnae |
| Caspase-13-like | Odobenus rosmarus divergens |
| Caspase-13-like-isoform-X1 | Balaenoptera acutorostrata scammoni |
| Caspase-13-like-isoform-X1 | Camelus bactrianus |
| Caspase-14 | Acinonyx jubatus |
| Caspase-14 | Alligator sinensis |
| Caspase-14 | Anolis carolinensis |
| Caspase-14 | Camelus bactrianus |
| Caspase-14 | Carlito syrichta |
| Caspase-14 | Chrysochloris asiatica |
| Caspase-14 | Equus caballus |
| Caspase-14 | Erinaceus europaeus |
| Caspase-14 | Odobenus rosmarus divergens |
| Caspase-14 | Python bivittatus |
| Caspase-14-like | Acinonyx jubatus |
| Caspase-14-like | Balaenoptera acutorostrata scammoni |
| Caspase-14-like | Camelus bactrianus |
| Caspase-14-like | Chelonia mydas |
| Caspase-14-like | Odobenus rosmarus divergens |
| Caspase-16 | Camelus bactrianus |
| Caspase-16 | Carlito syrichta |
| Caspase-16 | Pongo abelii |
| Caspase-16-isoform-X1 | Equus caballus |
| Caspase-17-A | Gallus gallus |
| Caspase-18-A | Gallus gallus |
| Caspase-1-A-like | Xenopus tropicalis |
| Caspase-1-isoform-X1 | Acinonyx jubatus |
| Caspase-1-isoform-X1 | Gallus gallus |
| Caspase-1-isoform-X1 | Pongo abelii |

| | |
|---|---|
| Caspase-1-like | Alligator sinensis |
| Caspase-1-like | Aptenodytes forsteri |
| Caspase-1-like | Apteryx australis mantelli |
| Caspase-1-like | Astyanax mexicanus |
| Caspase-1-like | Austrofundulus limnaeus |
| Caspase-1-like | Balaenoptera acutorostrata scammoni |
| Caspase-1-like | Callorhinchus milii |
| Caspase-1-like | Cariama cristata |
| Caspase-1-like | Carlito syrichta |
| Caspase-1-like | Chelonia mydas |
| Caspase-1-like | Chrysochloris asiatica |
| Caspase-1-like | Clupea harengus |
| Caspase-1-like | Gekko japonicus |
| Caspase-1-like | Nestor notabilis |
| Caspase-1-like | Python bivittatus |
| Caspase-1-like | Struthio camelus australis |
| Caspase-1-like | Tinamus guttatus |
| Caspase-1-like | Xiphophorus maculatus |
| Caspase-2 | Aptenodytes forsteri |
| Caspase-2 | Camelus bactrianus |
| Caspase-2 | Cariama cristata |
| Caspase-2 | Chelonia mydas |
| Caspase-2 | Chrysochloris asiatica |
| Caspase-2 | Clupea harengus |
| Caspase-2 | Danio rerio |
| Caspase-2 | Erinaceus europaeus |
| Caspase-2 | Gekko japonicus |
| Caspase-2 | Latimeria chalumnae |
| Caspase-2 | Melopsittacus undulatus |

| | |
|---|---|
| Caspase-2 | Nestor notabilis |
| Caspase-2 | Python bivittatus |
| Caspase-2 | Struthio camelus australis |
| Caspase-2 | Taeniopygia guttata |
| Caspase-2 | Xenopus tropicalis |
| Caspase-2 | Xiphophorus maculatus |
| Caspase-2-isoform-X1 | Acinonyx jubatus |
| Caspase-2-isoform-X1 | Alligator sinensis |
| Caspase-2-isoform-X1 | Anolis carolinensis |
| Caspase-2-isoform-X1 | Apteryx australis mantelli |
| Caspase-2-isoform-X1 | Astyanax mexicanus |
| Caspase-2-isoform-X1 | Austrofundulus limnaeus |
| Caspase-2-isoform-X1 | Balaenoptera acutorostrata scammoni |
| Caspase-2-isoform-X1 | Carlito syrichta |
| Caspase-2-isoform-X1 | Equus caballus |
| Caspase-2-isoform-X1 | Gallus gallus |
| Caspase-2-isoform-X1 | Odobenus rosmarus divergens |
| Caspase-2-isoform-X1 | Pongo abelii |
| Caspase-2-isoform-X1 | Tinamus guttatus |
| Caspase-3 | Alligator sinensis |
| Caspase-3 | Anolis carolinensis |
| Caspase-3 | Aptenodytes forsteri |
| Caspase-3 | Astyanax mexicanus |
| Caspase-3 | Balaenoptera acutorostrata scammoni |
| Caspase-3 | Callorhinchus milii |
| Caspase-3 | Camelus bactrianus |
| Caspase-3 | Cariama cristata |
| Caspase-3 | Carlito syrichta |
| Caspase-3 | Chelonia mydas |

| | |
|---|---|
| Caspase-3 | Chrysochloris asiatica |
| Caspase-3 | Clupea harengus |
| Caspase-3 | Danio rerio |
| Caspase-3 | Equus caballus |
| Caspase-3 | Erinaceus europaeus |
| Caspase-3 | Gallus gallus |
| Caspase-3 | Melopsittacus undulatus |
| Caspase-3 | Nestor notabilis |
| Caspase-3 | Pongo abelii |
| Caspase-3 | Python bivittatus |
| Caspase-3 | Xenopus tropicalis |
| Caspase-3 | Xiphophorus maculatus |
| Caspase-3-isoform-X1 | Apteryx australis mantelli |
| Caspase-3-isoform-X1 | Struthio camelus australis |
| Caspase-3-isoform-X3 | Odobenus rosmarus divergens |
| Caspase-3-isoform-X4 | Acinonyx jubatus |
| Caspase-3-like | Xiphophorus maculatus |
| Caspase-4-isoform-X1 | Equus caballus |
| Caspase-4-isoform-X1 | Pongo abelii |
| Caspase-4-like | Carlito syrichta |
| Caspase-5-isoform-X1 | Pongo abelii |
| Caspase-6 | Alligator sinensis |
| Caspase-6 | Anolis carolinensis |
| Caspase-6 | Aptenodytes forsteri |
| Caspase-6 | Austrofundulus limnaeus |
| Caspase-6 | Callorhinchus milii |
| Caspase-6 | Camelus bactrianus |
| Caspase-6 | Cariama cristata |
| Caspase-6 | Carlito syrichta |

| | |
|---|---|
| Caspase-6 | Chelonia mydas |
| Caspase-6 | Chrysochloris asiatica |
| Caspase-6 | Clupea harengus |
| Caspase-6 | Equus caballus |
| Caspase-6 | Erinaceus europaeus |
| Caspase-6 | Latimeria chalumnae |
| Caspase-6 | Melopsittacus undulatus |
| Caspase-6 | Nestor notabilis |
| Caspase-6 | Odobenus rosmarus divergens |
| Caspase-6 | Pongo abelii |
| Caspase-6 | Python bivittatus |
| Caspase-6 | Taeniopygia guttata |
| Caspase-6 | Xenopus tropicalis |
| Caspase-6 | Xiphophorus maculatus |
| Caspase-6-isoform-X1 | Apteryx australis mantelli |
| Caspase-6-isoform-X1 | Balaenoptera acutorostrata scammoni |
| Caspase-6-isoform-X1 | Gallus gallus |
| Caspase-6-isoform-X1 | Gekko japonicus |
| Caspase-6-isoform-X1 | Struthio camelus australis |
| Caspase-6-isoform-X2 | Acinonyx jubatus |
| Caspase-6-like-isoform-X1 | Danio rerio |
| Caspase-7 | Acinonyx jubatus |
| Caspase-7 | Aptenodytes forsteri |
| Caspase-7 | Apteryx australis mantelli |
| Caspase-7 | Astyanax mexicanus |
| Caspase-7 | Camelus bactrianus |
| Caspase-7 | Cariama cristata |
| Caspase-7 | Chrysochloris asiatica |
| Caspase-7 | Clupea harengus |

| | |
|---|---|
| Caspase-7 | Danio rerio |
| Caspase-7 | Equus caballus |
| Caspase-7 | Erinaceus europaeus |
| Caspase-7 | Gallus gallus |
| Caspase-7 | Nestor notabilis |
| Caspase-7 | Pongo abelii |
| Caspase-7 | Python bivittatus |
| Caspase-7 | Struthio camelus australis |
| Caspase-7 | Taeniopygia guttata |
| Caspase-7 | Tinamus guttatus |
| Caspase-7 | Xenopus tropicalis |
| Caspase-7 | Xiphophorus maculatus |
| Caspase-7-isoform-X1 | Anolis carolinensis |
| Caspase-7-isoform-X1 | Austrofundulus limnaeus |
| Caspase-7-isoform-X1 | Balaenoptera acutorostrata scammoni |
| Caspase-7-isoform-X1 | Carlito syrichta |
| Caspase-7-isoform-X1 | Latimeria chalumnae |
| Caspase-7-isoform-X1 | Melopsittacus undulatus |
| Caspase-7-isoform-X1 | Odobenus rosmarus divergens |
| Caspase-7-isoform-X2 | Alligator sinensis |
| Caspase-7-like | Callorhinchus milii |
| Caspase-7-like | Gekko japonicus |
| Caspase-8 | Acinonyx jubatus |
| Caspase-8 | Aptenodytes forsteri |
| Caspase-8 | Apteryx australis mantelli |
| Caspase-8 | Callorhinchus milii |
| Caspase-8 | Cariama cristata |
| Caspase-8 | Carlito syrichta |
| Caspase-8 | Danio rerio |

| | |
|---|---|
| Caspase-8 | Equus caballus |
| Caspase-8 | Erinaceus europaeus |
| Caspase-8 | Nestor notabilis |
| Caspase-8 | Pongo abelii |
| Caspase-8 | Struthio camelus australis |
| Caspase-8 | Xenopus tropicalis |
| Caspase-8-isoform-X1 | Balaenoptera acutorostrata scammoni |
| Caspase-8-isoform-X1 | Camelus bactrianus |
| Caspase-8-isoform-X1 | Chrysochloris asiatica |
| Caspase-8-isoform-X1 | Odobenus rosmarus divergens |
| Caspase-8-like | Alligator sinensis |
| Caspase-8-like | Aptenodytes forsteri |
| Caspase-8-like | Apteryx australis mantelli |
| Caspase-8-like | Astyanax mexicanus |
| Caspase-8-like | Austrofundulus limnaeus |
| Caspase-8-like | Chelonia mydas |
| Caspase-8-like | Clupea harengus |
| Caspase-8-like | Latimeria chalumnae |
| Caspase-8-like | Melopsittacus undulatus |
| Caspase-8-like | Nestor notabilis |
| Caspase-8-like | Struthio camelus australis |
| Caspase-8-like | Taeniopygia guttata |
| Caspase-8-like | Tinamus guttatus |
| Caspase-8-like | Xiphophorus maculatus |
| Caspase-8-like-isoform-X1 | Astyanax mexicanus |
| Caspase-8-like-isoform-X1 | Melopsittacus undulatus |
| Caspase-9 | Alligator sinensis |
| Caspase-9 | Anolis carolinensis |
| Caspase-9 | Aptenodytes forsteri |

| | |
|---|---|
| Caspase-9 | Apteryx australis mantelli |
| Caspase-9 | Austrofundulus limnaeus |
| Caspase-9 | Callorhinchus milii |
| Caspase-9 | Camelus bactrianus |
| Caspase-9 | Carlito syrichta |
| Caspase-9 | Chelonia mydas |
| Caspase-9 | Chrysochloris asiatica |
| Caspase-9 | Clupea harengus |
| Caspase-9 | Danio rerio |
| Caspase-9 | Gallus gallus |
| Caspase-9 | Gekko japonicus |
| Caspase-9 | Melopsittacus undulatus |
| Caspase-9 | Pongo abelii |
| Caspase-9 | Struthio camelus australis |
| Caspase-9 | Taeniopygia guttata |
| Caspase-9 | Tinamus guttatus |
| Caspase-9 | Xenopus tropicalis |
| Caspase-9 | Xiphophorus maculatus |
| Caspase-9-isoform-X1 | Acinonyx jubatus |
| Caspase-9-isoform-X1 | Erinaceus europaeus |
| Caspase-9-isoform-X1 | Odobenus rosmarus divergens |
| Caspase-Xa | Danio rerio |
| Caspy2 | Danio rerio |

**Supplemental Table S2. APR_2: List of Taxa and caspases used in reconstructions from the CaspBase.**

| Caspase | Species |
|---|---|
| Caspase-1 | Camelus bactrianus |
| Caspase-1 | Canis lupus familiaris |

| | |
|---|---|
| Caspase-1 | Columba livia |
| Caspase-1 | Danio rerio |
| Caspase-1 | Dasypus novemcinctus |
| Caspase-1 | Homo sapiens |
| Caspase-1 | Lepisosteus oculatus |
| Caspase-1 | Mus musculus |
| Caspase-1 | Myotis brandtii |
| Caspase-1 | Orycteropus afer afer |
| Caspase-1 | Rattus norvegicus |
| Caspase-1 | Sus scrofa |
| Caspase-10 | Ailuropoda melanol |
| Caspase-10 | Alligator mississippiensis |
| Caspase-10 | Anas platyrhynchos |
| Caspase-10 | Aptenodytes forsteri |
| Caspase-10 | Canis lupus familiaris |
| Caspase-10 | Dasypus novemcinctus |
| Caspase-10 | Felis catusdomestic |
| Caspase-10 | Gallus gallus |
| Caspase-10 | Homo sapiens |
| Caspase-10 | Sarcophilus harrisii |
| Caspase-10 | Sus scrofa |
| Caspase-10 | Xenopus tropicalis |
| Caspase-10-like | Gekko japonicus |
| Caspase-10-like | Struthio camelus australis |
| Caspase-11 | Rattus norvegicus |
| Caspase-13 | Bison bison bison |
| Caspase-13 | Dasypus novemcinctus |
| Caspase-13 | Sus scrofa |
| Caspase-13-like | Camelus bactrianus |

| | |
|---|---|
| Caspase-13-like | Orycteropus afer afer |
| Caspase-13-like | Physeter catodon |
| Caspase-14 | Ailuropoda melanol |
| Caspase-14 | Bison bison bison |
| Caspase-14 | Camelus bactrianus |
| Caspase-14 | Dasypus novemcinctus |
| Caspase-14 | Felis catusdomestic |
| Caspase-14 | Homo sapiens |
| Caspase-14 | Myotis brandtii 1 |
| Caspase-14 | Myotis brandtii 2 |
| Caspase-14 | Orycteropus afer afer |
| Caspase-14 | Rattus norvegicus |
| Caspase-14 | Sarcophilus harrisii |
| Caspase-14 | Sus scrofa |
| Caspase-14-like | Sus scrofa |
| Caspase-15 | Sus scrofa |
| Caspase-18-A | Gallus gallus |
| Caspase-1-A | Gallus gallus |
| Caspase-1-b | Gallus gallus |
| Caspase-1-like | Alligator mississippiensis |
| Caspase-1-like | Aptenodytes forsteri |
| Caspase-1-like | Bison bison bison |
| Caspase-1-like | Callorhinchus milii |
| Caspase-1-like | Canis lupus familiaris |
| Caspase-1-like | Gekko japonicus |
| Caspase-1-like | Myotis brandtii |
| Caspase-1-like | Physeter catodon |
| Caspase-1-like | Python bivittatus |
| Caspase-1-like | Sarcophilus harrisii |

| | |
|---|---|
| Caspase-1-like | Sarcophilus harrisii |
| Caspase-1-like | Stegastes partitus |
| Caspase-1-like | Struthio camelus australis |
| Caspase-2 | Ailuropoda melanol |
| Caspase-2 | Alligator mississippiensis |
| Caspase-2 | Anas platyrhynchos |
| Caspase-2 | Anolis carolinensis |
| Caspase-2 | Aptenodytes forsteri |
| Caspase-2 | Bison bison bison |
| Caspase-2 | Camelus bactrianus |
| Caspase-2 | Canis lupus familiaris |
| Caspase-2 | Columba livia |
| Caspase-2 | Dasypus novemcinctus |
| Caspase-2 | Felis catusdomestic |
| Caspase-2 | Gekko japonicus |
| Caspase-2 | Homo sapiens |
| Caspase-2 | Latimeria chalumnae |
| Caspase-2 | Mus musculus |
| Caspase-2 | Myotis brandtii |
| Caspase-2 | Ornithorhynchus an |
| Caspase-2 | Orycteropus afer afer |
| Caspase-2 | Physeter catodon |
| Caspase-2 | Python bivittatus |
| Caspase-2 | Rattus norvegicus |
| Caspase-2 | Sarcophilus harrisii |
| Caspase-2 | Struthio camelus australis |
| Caspase-2 | Xenopus tropicalis |
| Caspase-2-A | Gallus gallus |
| Caspase-2-B | Gallus gallus |

| | |
|---|---|
| Caspase-2-like | Sarcophilus harrisii |
| Caspase-3 | Ailuropoda melanol |
| Caspase-3 | Alligator mississippiensis |
| Caspase-3 | Anas platyrhynchos |
| Caspase-3 | Aptenodytes forsteri |
| Caspase-3 | Bison bison bison |
| Caspase-3 | Callorhinchus milii |
| Caspase-3 | Camelus bactrianus |
| Caspase-3 | Canis lupus familiaris |
| Caspase-3 | Columba livia |
| Caspase-3 | Cynoglossus semilaevis |
| Caspase-3 | Dasypus novemcinctus |
| Caspase-3 | Felis catusdomestic |
| Caspase-3 | Gallus gallus |
| Caspase-3 | Homo sapiens |
| Caspase-3 | Mus musculus |
| Caspase-3 | Myotis brandtii |
| Caspase-3 | Ornithorhynchus anatinus |
| Caspase-3 | Orycteropus afer afer |
| Caspase-3 | Physeter catodon |
| Caspase-3 | Python bivittatus |
| Caspase-3 | Rattus norvegicus |
| Caspase-3 | Sarcophilus harrisii |
| Caspase-3 | Sarcophilus harrisii |
| Caspase-3 | Struthio camelus australis |
| Caspase-3 | Sus scrofa |
| Caspase-3 | Takifugu rubripes |
| Caspase-3 | Xenopus tropicalis |
| Caspase-3-A | Anolis carolinensis |

| | |
|---|---|
| Caspase-3-A | Danio rerio |
| Caspase-3-A | Oryzias latipes |
| Caspase-3-B | Danio rerio |
| Caspase-3-B | Oryzias latipes |
| Caspase-3-like | Austrofundulus limnaeus |
| Caspase-3-like | Cynoglossus semilaevis |
| Caspase-3-like-A | Stegastes partitus |
| Caspase-3-like-C | Stegastes partitus |
| Caspase-4 | Homo sapiens |
| Caspase-5 | Homo sapiens |
| Caspase-6 | Ailuropoda melanol |
| Caspase-6 | Alligator mississippiensis |
| Caspase-6 | Anas platyrhynchos |
| Caspase-6 | Anolis carolinensis |
| Caspase-6 | Aptenodytes forsteri |
| Caspase-6 | Austrofundulus limnaeus |
| Caspase-6 | Bison bison bison |
| Caspase-6 | Callorhinchus milii |
| Caspase-6 | Camelus bactrianus |
| Caspase-6 | Canis lupus familiaris |
| Caspase-6 | Columba livia |
| Caspase-6 | Cynoglossus semilaevis |
| Caspase-6 | Danio rerio |
| Caspase-6 | Dasypus novemcinctus |
| Caspase-6 | Felis catusdomestic |
| Caspase-6 | Gallus gallus |
| Caspase-6 | Gallus gallus 2 |
| Caspase-6 | Gekko japonicus |
| Caspase-6 | Latimeria chalumnae |

| | |
|---|---|
| Caspase-6 | Lepisosteus oculatus |
| Caspase-6 | Mus musculus |
| Caspase-6 | Myotis brandtii |
| Caspase-6 | Ornithorhynchus anatinus |
| Caspase-6 | Orycteropus afer afer |
| Caspase-6 | Oryzias latipes |
| Caspase-6 | Physeter catodon |
| Caspase-6 | Python bivittatus |
| Caspase-6 | Rattus norvegicus |
| Caspase-6 | Sarcophilus harrisii |
| Caspase-6 | Stegastes partitus |
| Caspase-6 | Struthio camelus australis |
| Caspase-6 | Sus scrofa |
| Caspase-6 | Takifugu rubripes |
| Caspase-6 | Xenopus tropicalis |
| Caspase-6-like | Danio rerio |
| Caspase-7 | Ailuropoda melanol |
| Caspase-7 | Alligator mississippiensis |
| Caspase-7 | Anas platyrhynchos |
| Caspase-7 | Anolis carolinensis |
| Caspase-7 | Aptenodytes forsteri |
| Caspase-7 | Bison bison bison |
| Caspase-7 | Camelus bactrianus |
| Caspase-7 | Canis lupus familiaris |
| Caspase-7 | Columba livia |
| Caspase-7 | Cynoglossus semilaevis |
| Caspase-7 | Danio rerio |
| Caspase-7 | Dasypus novemcinctus |
| Caspase-7 | Felis catusdomestic |

| | |
|---|---|
| Caspase-7 | Gallus gallus |
| Caspase-7 | Homo sapiens |
| Caspase-7 | Latimeria chalumnae |
| Caspase-7 | Lepisosteus oculatus |
| Caspase-7 | Mus musculus |
| Caspase-7 | Ornithorhynchus anatinus |
| Caspase-7 | Physeter catodon |
| Caspase-7 | Python bivittatus |
| Caspase-7 | Rattus norvegicus |
| Caspase-7 | Sarcophilus harrisii |
| Caspase-7 | Sarcophilus harrisii |
| Caspase-7 | Stegastes partitus |
| Caspase-7 | Struthio camelus australis |
| Caspase-7 | Takifugu rubripes |
| Caspase-7 | Xenopus tropicalis |
| Caspase-7-like | Callorhinchus milii |
| Caspase-7-like-A | Gekko japonicus |
| Caspase-7-like-B | Gekko japonicus |
| Caspase-8 | Ailuropoda melanol |
| Caspase-8 | Anas platyrhynchos |
| Caspase-8 | Aptenodytes forsteri |
| Caspase-8 | Bison bison bison |
| Caspase-8 | Camelus bactrianus |
| Caspase-8 | Canis lupus familiaris |
| Caspase-8 | Dasypus novemcinctus |
| Caspase-8 | Felis catusdomestic |
| Caspase-8 | Gallus gallus |
| Caspase-8 | Homo sapiens |
| Caspase-8 | Lepisosteus oculatus |

| | |
|---|---|
| Caspase-8 | Mus musculus |
| Caspase-8 | Myotis brandtii |
| Caspase-8 | Orycteropus afer afer |
| Caspase-8 | Physeter catodon |
| Caspase-8 | Rattus norvegicus |
| Caspase-8 | Struthio camelus australis |
| Caspase-8 | Sus scrofa |
| Caspase-8 | Xenopus tropicalis |
| Caspase-8-B | Danio rerio |
| Caspase-8-like | Alligator mississippiensis |
| Caspase-8-like | Anas platyrhynchos |
| Caspase-8-like | Aptenodytes forsteri |
| Caspase-8-like | Austrofundulus limnaeus |
| Caspase-8-like | Struthio camelus australis |
| Caspase-8-like-A | Latimeria chalumnae |
| Caspase-8-like-A | Lepisosteus oculatus |
| Caspase-8-like-A | Stegastes partitus |
| Caspase-8-like-A | Takifugu rubripes |
| Caspase-8-like-B | Latimeria chalumnae |
| Caspase-8-like-B | Lepisosteus oculatus |
| Caspase-9 | Ailuropoda melanol |
| Caspase-9 | Alligator mississippiensis |
| Caspase-9 | Anas platyrhynchos |
| Caspase-9 | Anolis carolinensis |
| Caspase-9 | Aptenodytes forsteri |
| Caspase-9 | Austrofundulus limnaeus |
| Caspase-9 | Bison bison bison |
| Caspase-9 | Callorhinchus milii |
| Caspase-9 | Camelus bactrianus |

| | |
|---|---|
| Caspase-9 | Canis lupus familiaris |
| Caspase-9 | Columba livia |
| Caspase-9 | Cynoglossus semilaevis |
| Caspase-9 | Danio rerio |
| Caspase-9 | Dasypus novemcinctus |
| Caspase-9 | Felis catusdomestic |
| Caspase-9 | Gallus gallus |
| Caspase-9 | Gekko japonicus |
| Caspase-9 | Homo sapiens |
| Caspase-9 | Lepisosteus oculatus |
| Caspase-9 | Mus musculus |
| Caspase-9 | Myotis brandtii |
| Caspase-9 | Ornithorhynchus anatinus |
| Caspase-9 | Orycteropus afer afer |
| Caspase-9 | Physeter catodon |
| Caspase-9 | Rattus norvegicus |
| Caspase-9 | Sarcophilus harrisii |
| Caspase-9 | Sarcophilus harrisii |
| Caspase-9 | Stegastes partitus |
| Caspase-9 | Struthio camelus australis |
| Caspase-9 | Sus scrofa |
| Caspase-9 | Takifugu rubripes |
| Caspase-9-like | Sarcophilus harrisii |
| initiator | Alligator mississippiensis |

**Supplemental Table S3. Characteristics of ancestral caspases.**

| Composition[1] | Protein | | | | | |
|---|---|---|---|---|---|---|
| | AncCP-Ef1 | AncCP-Ef2 | AncCP-6An | Casp-3 | Casp-6 | Casp-7 |
| Total Number Amino Acids | 277 | 275 | 290 | 277 | 293 | 303 |
| Ala (A) | 10 (3.6%) | 9 (3.3%) | 22 (7.6%) | 12 (4.3%) | 19 (6.5%) | 18 (5.9%) |
| Arg (R) | 8 (2.9%) | 7 (2.5%) | 15 (5.2%) | 14 (5.1%) | 17 (5.8%) | 15 (5.0%) |
| Asn (N) | 18 (6.5%) | 13 (4.7%) | 13 (4.5%) | 15 (5.4%) | 11 (3.8%) | 14 (4.6%) |
| Asp (D) | 19 (6.9%) | 20 (7.3%) | 22 (7.6%) | 20 (7.2%) | 20 (6.8%) | 27 (8.9%) |
| Cys (C) | 7 (2.5%) | 7 (2.5%) | 8 (2.8%) | 8 (2.9%) | 10 (3.4%) | 11 (3.6%) |
| Gln (Q) | 7 (2.5%) | 10 (3.6%) | 8 (2.8%) | 4 (1.4%) | 7 (2.4%) | 11 (3.6%) |
| Glu (E) | 24 (8.7%) | 25 (9.1%) | 26 (9.0%) | 20 (7.2%) | 20 (6.8%) | 19 (6.3%) |
| Gly (G) | 20 (7.2%) | 18 (6.5%) | 19 (6.6%) | 16 (5.8%) | 19 (6.5%) | 18 (5.9%) |
| His (H) | 6 (2.2%) | 6 (2.2%) | 5 (1.7%) | 8 (2.9%) | 12 (4.1%) | 7 (2.3%) |
| Ile (I) | 14 (5.1%) | 14 (5.1%) | 11 (3.8%) | 19 (6.9%) | 13 (4.4%) | 17 (5.6%) |
| Leu (L) | 24 (8.7%) | 25 (9.1%) | 23 (7.9%) | 20 (7.2%) | 26 (8.9%) | 20 (6.6%) |
| Lys (K) | 25 (9.0%) | 28 (10.2%) | 25 (8.6%) | 22 (7.9%) | 20 (6.8%) | 25 (8.3%) |
| Met (M) | 8 (2.9%) | 6 (2.2%) | 10 (3.4%) | 10 (3.6%) | 7 (2.4%) | 7 (2.3%) |
| Phe (F) | 12 (4.3%) | 12 (4.4%) | 14 (4.8%) | 15 (5.4%) | 18 (6.1%) | 17 (5.6%) |
| Pro (P) | 8 (2.9%) | 9 (3.3%) | 9 (3.1%) | 7 (2.5%) | 10 (3.4%) | 12 (4.0%) |
| Ser (S) | 30 (10.8%) | 26 (9.5%) | 16 (5.5%) | 26 (9.4%) | 18 (6.1%) | 21 (6.9%) |
| Thr (T) | 13 (4.7%) | 13 (4.7%) | 15 (5.2%) | 16 (5.8%) | 16 (5.5%) | 15 (5.0%) |
| Trp (W) | 1 (0.4%) | 1 (0.4%) | 2 (0.7%) | 2 (0.7%) | 2 (0.7%) | 2 (0.7%) |
| Tyr (Y) | 9 (3.2%) | 13 (4.7%) | 12 (4.1%) | 10 (3.6%) | 10 (3.4%) | 9 (3.0%) |
| Val (V) | 14 (5.1%) | 13 (4.7%) | 15 (5.2%) | 13 (4.7%) | 18 (6.1%) | 18 (5.9%) |

| | | | | | | |
|---|---|---|---|---|---|---|
| Molecular Weight (Da) | 30,967 | 31,221 | 32,942 | 31,608 | 33,310 | 34,277 |
| pI | 5.22 | 5.20 | 5.35 | 6.09 | 6.46 | 5.72 |
| Extinction Coefficient (280 nm, $M^{-1}cm^{-1}$)[(2)] | 18,910 | 24,870 | 28,880 | 25,900 | 25,900 | 24,410 |

[1] Parameters exclude the LEHHHHHH C-terminal tag.

[2] Assuming all cysteine residues are reduced.

**Supplemental Table S4. Crystallographic parameters for ancestral caspases.**

| | **AncCP-Ef1 (DEVD)** | **AncCP-6An (VEID)** |
|---|---|---|
| PDB Code | 6PDQ | 6PPM |
| Data collection | | |
| Wavelength (Å) | 1.0 | 1.0 |
| Temperature (K) | 100 | 100 |
| Space Group | $P2_12_12_1$ | $P2_12_12_1$ |
| Cell Dimensions | | |
| a, b, c (Å) | 52.17, 78.76, 107.74 | 84.31, 88.54, 141.03 |
| α, β, γ (°) | 90, 90, 90 | 90, 90, 90 |
| # Unique Reflections | 39730 (3741) | 32724 (3134) |
| Resolution (Å) | 43.49-1.83 (1.89-1.83) | 42.24-2.61 (2.70-2.61) |
| R-meas | 0.113 (0.620) | 0.171 (1.510) |
| R-pim | 0.041 (0.227) | 0.063 (0.547) |
| CC (1/2) | 0.977 (0.914) | 0.951 (0.827) |
| Average I/σ | 8.5 (2.6) | 5.1 (2.4) |
| Completeness (%) | 99.23 (95.28) | 99.68 (97.69) |
| Redundancy | 7.0 (5.3) | 3.7 |
| Refinement | | |
| $R_{work}$ / $R_{free}$ | 0.192 / 0.235 | 0.178 / 0.251 |
| Average B-factor ($Å^2$) | 32.85 | 54.38 |
| Macromolecules | 32.69 | 54.60 |
| Solvent | 36.14 | 49.40 |
| Wilson B-factor | 28.56 | 51.80 |
| R. m. s. deviations | | |
| bond lengths (Å) | 0.006 | 0.007 |
| bond angles (°) | 0.81 | 0.91 |
| MolProbity Score | 1.25 | 1.70 |
| Number of atoms | | |

| | | |
|---|---|---|
| Protein | 3547 | 7639 |
| Water | 147 | 34 |
| Protein Residues | 463 | 985 |
| Clashscore | 2.9 | 7.02 |
| MolProbity | | |
| Ramachandran favored (%) | 98.2 | 95.46 |
| Ramachandran outliers (%) | 0.0 | 0.11 |
| Rotamer Outliers | 0.53 | 0.39 |

**Supplemental Table S5. Summary of enzyme activity.**

| Protein | Substrate | $k_{cat}$ ($sec^{-1}$)[(1)] | $K_M$ (μM) | $k_{cat}/K_M$ ($M^{-1}sec^{-1}$) |
|---|---|---|---|---|
| AncCP-Ef1 | VEID | 0.3 | 91 | $3.3x10^3$ |
| AncCP-Ef1 | LETD | 0.3 | 89 | $3.4x10^3$ |
| AncCP-Ef1 | DEVD | 0.8 | 117 | $6.8x10^3$ |
| AncCP-EF2 | VEID | ND | >300 | ND |
| AncCP-Ef2 | DEVD | 0.7 | 246 | $2.8x10^3$ |
| AncCP-Ef1 (CP-N162E) | VEID | 0.5 | 42 | $1.2x10^4$ |
| AncCP-Ef1 (CP-N162E) | LETD | 0.9 | 79 | $1.3x10^4$ |
| AncCP-Ef1 (CP-N162E) | DEVD | 3.4 | 250 | $1.4x10^4$ |
| AncCP-Ef1(E248V) | VEID | 0.04 | 195 | $2.1x10^2$ |
| AncCP-Ef1(E248V) | DEVD | 0.15 | 439 | $3.4x10^2$ |
| AncCP-Ef1-DM[(2)] | VEID | 1.1 | 32 | $3.4x10^4$ |
| AncCP-Ef1-DM | LETD | 1.9 | 45 | $4.2x10^4$ |
| AncCP-Ef1-DM | DEVD | 4.0 | 145 | $2.8x10^4$ |
| AncCP-Ef1-DM -3M[(3)] | VEID | 0.4 | 1.5 | $2.7x10^5$ |
| AncCP-Ef1-DM -3M | LETD | 1.2 | 19 | $6.3x10^4$ |
| AncCP-Ef1-DM -3M | DEVD | 2.1 | 55 | $3.8x10^4$ |
| AncCP-Ef1-DM -5M[(4)] | VEID | 0.6 | 4 | $1.5x10^5$ |

| | | | | |
|---|---|---|---|---|
| AncCP-Ef1-DM -5M | LETD | 1.5 | 20 | $7.5x10^4$ |
| AncCP-Ef1-DM -5M | DEVD | 2.9 | 150 | $1.9x10^4$ |
| AncCP-6An | VEID | 0.24 | 25 | $9.6x10^3$ |
| AncCP-6An | LETD | 1.2 | 300 | $4x10^3$ |
| AncCP-6An | DEVD | ND | >500 | ND |
| HsCaspase-6 | VEID | 1.2 | 37 | $3.2x10^4$ |
| HsCaspase-6 | DEVD | 0.6 | 460 | $1.3x10^3$ |

[1] Standard error for all measurements was <10%.

[2] AncCP-Ef1(DM), Ancestral effector caspase, first resurrection (AncCP-Ef1) with CP-N162E and GP9-E01V mutation.

[3] AncCP-Ef1(3M), AncCP-Ef1(DM) + CP-S172D

[4] AncCP-Ef1(5M), AncCP-Ef1(3M) + CP-S199R + GP9-S02D

### Preliminary Full wwPDB X-ray Structure Validation Report (i)

Jun 19, 2019 – 12:46 PM EDT

| | | |
|---|---|---|
| Deposition ID | : | D_1000242366 |
| PDB ID | : | *(not yet assigned)* |

This is a Preliminary Full wwPDB X-ray Structure Validation Report.

This report is produced by the wwPDB Deposition System during initial deposition but before annotation of the structure.

We welcome your comments at **
A user guide is available at
https://www.wwpdb.org/validation/2017/XrayValidationReportHelp
with specific help available everywhere you see the (i) symbol.

PRELIMINARY VALIDATION REPORT

The following versions of software and data (see references (i)) were used in the production of this report:

| | | |
|---|---|---|
| MolProbity | : | 4.02b-467 |
| Mogul | : | 1.8.0 (224370), CSD as540be (2019) |
| Xtriage (Phenix) | : | 1.13 |
| EDS | : | 2.3.2 |
| Percentile statistics | : | 20171227.v01 (using entries in the PDB archive December 27th 2017) |
| Refmac | : | 5.8.0158 |
| CCP4 | : | 7.0 (Gargrove) |
| Ideal geometry (proteins) | : | Engh & Huber (2001) |
| Ideal geometry (DNA, RNA) | : | Parkinson et al. (1996) |
| Validation Pipeline (wwPDB-VP) | : | 2.3.2 |

### 1 Overall quality at a glance (i)

The following experimental techniques were used to determine the structure:
*X-RAY DIFFRACTION*

The reported resolution of this entry is 1.83 Å.

Percentile scores (ranging between 0-100) for global validation metrics of the entry are shown in the following graphic. The table shows the number of entries on which the scores are based.

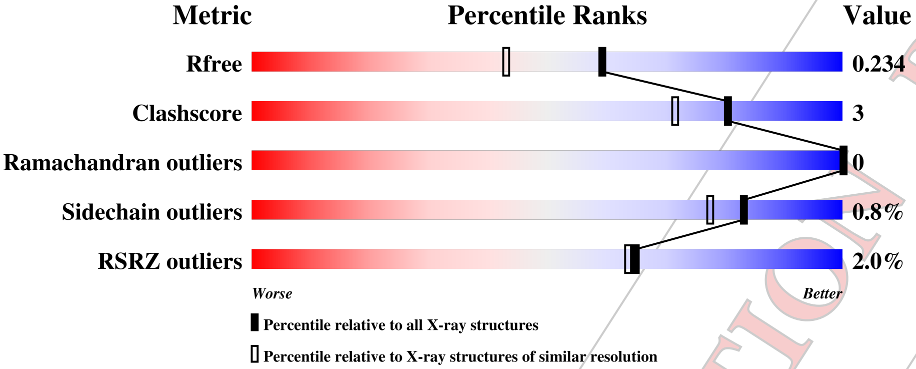


| Metric | Whole archive (#Entries) | Similar resolution (#Entries, resolution range(Å)) |
|---|---|---|
| $R_{free}$ | 111664 | 3313 (1.86-1.82) |
| Clashscore | 122126 | 3530 (1.86-1.82) |
| Ramachandran outliers | 120053 | 3495 (1.86-1.82) |
| Sidechain outliers | 120020 | 3496 (1.86-1.82) |
| RSRZ outliers | 108989 | 3265 (1.86-1.82) |

The table below summarises the geometric issues observed across the polymeric chains and their fit to the electron density. The red, orange, yellow and green segments on the lower bar indicate the fraction of residues that contain outliers for >=3, 2, 1 and 0 types of geometric quality criteria. A grey segment represents the fraction of residues that are not modelled. The numeric value for each fraction is indicated below the corresponding segment, with a dot representing fractions <=5% The upper red bar (where present) indicates the fraction of residues that have poor fit to the electron density. The numeric value is given above the bar.

| Mol | Chain | Length | Quality of chain |
|---|---|---|---|
| 1 | A | 245 | % / 54% • 43% |
| 1 | B | 245 | 33% • 64% |
| 2 | D | 142 | 3% / 94% 6% • |
| 3 | E | 85 | 4% / 89% 9% • |
| 4 | F | 5 | 60% 40% |

*Continued on next page...*

PRELIMINARY VALIDATION REPORT

*Continued from previous page...*

| Mol | Chain | Length | Quality of chain |
|---|---|---|---|
| 4 | G | 5 | 100% |

PRELIMINARY VALIDATION REPORT

#### 2 Entry composition (i)

There are 5 unique types of molecules in this entry. The entry contains 3756 atoms, of which 0 are hydrogens and 0 are deuteriums.

In the tables below, the ZeroOcc column contains the number of atoms modelled with zero occupancy, the AltConf column contains the number of residues with at least one atom in alternate conformation and the Trace column contains the number of residues modelled with at most 2 atoms.

- Molecule 1 is a protein called Ancestral Effector Caspase-3/6/7.

| Mol | Chain | Residues | Atoms | | | | | ZeroOcc | AltConf | Trace |
|---|---|---|---|---|---|---|---|---|---|---|
| 1 | A | 139 | Total | C | N | O | S | 0 | 3 | 0 |
| | | | 1063 | 666 | 178 | 211 | 8 | | | |
| 1 | B | 87 | Total | C | N | O | S | 0 | 2 | 0 |
| | | | 686 | 447 | 107 | 125 | 7 | | | |

- Molecule 2 is a protein.

| Mol | Chain | Residues | Atoms | | | | | ZeroOcc | AltConf | Trace |
|---|---|---|---|---|---|---|---|---|---|---|
| 2 | D | 142 | Total | C | N | O | S | 0 | 2 | 0 |
| | | | 1085 | 678 | 184 | 215 | 8 | | | |

- Molecule 3 is a protein.

| Mol | Chain | Residues | Atoms | | | | | ZeroOcc | AltConf | Trace |
|---|---|---|---|---|---|---|---|---|---|---|
| 3 | E | 85 | Total | C | N | O | S | 0 | 2 | 0 |
| | | | 673 | 438 | 106 | 123 | 6 | | | |

- Molecule 4 is a protein called ACE-ASP-GLU-VAL-ASP.

| Mol | Chain | Residues | Atoms | | | | ZeroOcc | AltConf | Trace |
|---|---|---|---|---|---|---|---|---|---|
| 4 | F | 5 | Total | C | N | O | 0 | 0 | 0 |
| | | | 35 | 20 | 4 | 11 | | | |
| 4 | G | 5 | Total | C | N | O | 0 | 0 | 0 |
| | | | 35 | 20 | 4 | 11 | | | |

- Molecule 5 is water.

| Mol | Chain | Residues | Atoms | | ZeroOcc | AltConf |
|---|---|---|---|---|---|---|
| 5 | C | 179 | Total | O | 0 | 0 |
| | | | 179 | 179 | | |

PRELIMINARY VALIDATION REPORT

### 3 Residue-property plots (i)

These plots are drawn for all protein, RNA and DNA chains in the entry. The first graphic for a chain summarises the proportions of the various outlier classes displayed in the second graphic. The second graphic shows the sequence view annotated by issues in geometryand electron density. Residues are color-coded according to the number of geometric quality criteria for which they contain at least one outlier: green = 0, yellow = 1, orange = 2 and red = 3 or more. A red dot above a residue indicates a poor fit to the electron density (RSRZ > 2). Stretches of 2 or more consecutive residues without any outlier are shown as a green connector. Residues present in the sample, but not in the model, are shown in grey.

- Molecule 1: Ancestral Effector Caspase-3/6/7

Chain A:

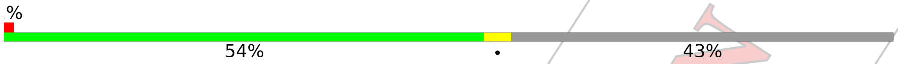


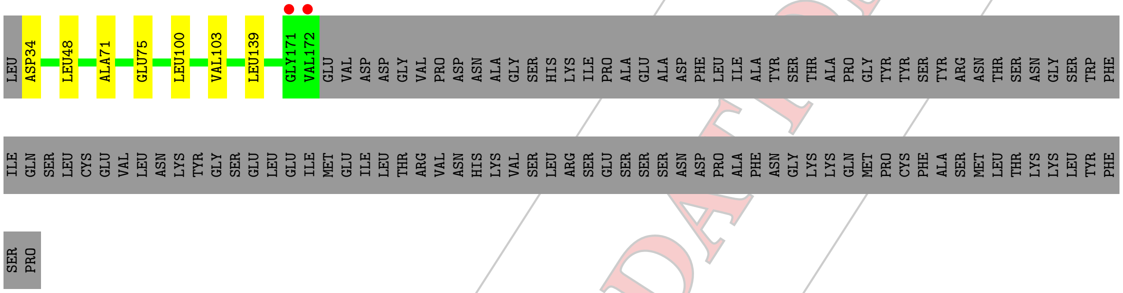


- Molecule 1: Ancestral Effector Caspase-3/6/7

Chain B:

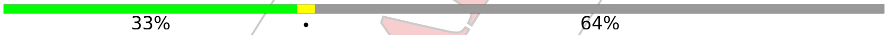


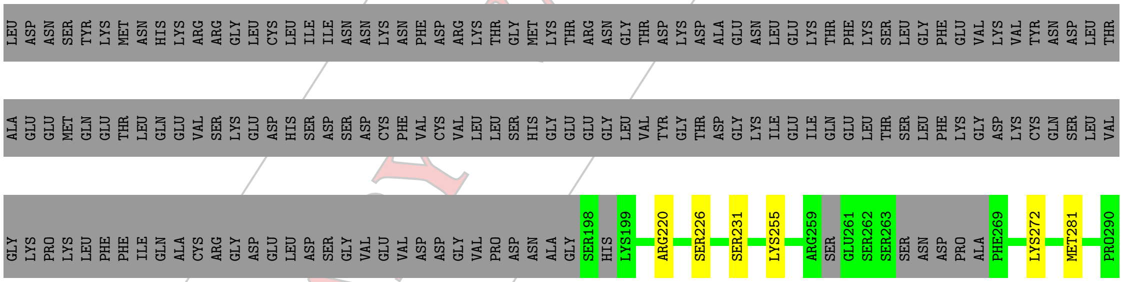


- Molecule 2:

Chain D:

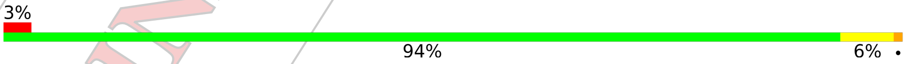


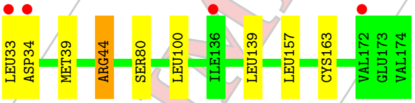


- Molecule 3:

Chain E:

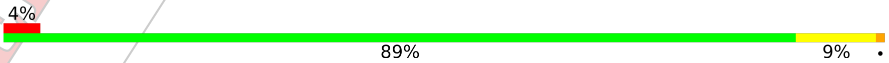


PRELIMINARY VALIDATION REPORT

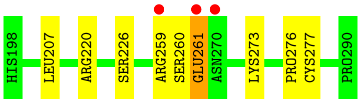


• Molecule 4: ACE-ASP-GLU-VAL-ASP

Chain F:
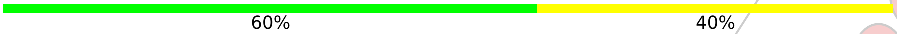


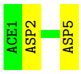


• Molecule 4: ACE-ASP-GLU-VAL-ASP

Chain G:
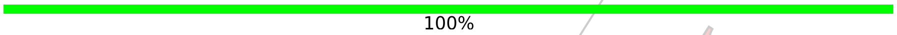


There are no outlier residues recorded for this chain.

PRELIMINARY VALIDATION REPORT

### 4 Data and refinement statistics (i)

| Property | Value | Source |
|---|---|---|
| Space group | P 21 21 21 | Depositor |
| Cell constants<br>a, b, c, $\alpha$, $\beta$, $\gamma$ | 52.17Å 78.76Å 107.74Å<br>90.00° 90.00° 90.00° | Depositor |
| Resolution (Å) | 43.49 – 1.83<br>43.49 – 1.83 | Depositor<br>EDS |
| % Data completeness<br>(in resolution range) | 99.2 (43.49-1.83)<br>99.2 (43.49-1.83) | Depositor<br>EDS |
| $R_{merge}$ | (Not available) | Depositor |
| $R_{sym}$ | (Not available) | Depositor |
| $< I/\sigma(I) >$ [1] | 1.78 (at 1.83Å) | Xtriage |
| Refinement program | PHENIX (1.15.2_3472: ???) | Depositor |
| R, $R_{free}$ | 0.191 , 0.234<br>0.191 , 0.234 | Depositor<br>DCC |
| $R_{free}$ test set | 1993 reflections (5.03%) | wwPDB-VP |
| Wilson B-factor (Å$^2$) | 28.6 | Xtriage |
| Anisotropy | 0.549 | Xtriage |
| Bulk solvent $k_{sol}$(e/Å$^3$), $B_{sol}$(Å$^2$) | 0.32 , 46.5 | EDS |
| L-test for twinning[2] | $< \|L\| > = 0.49$, $< L^2 > = 0.33$ | Xtriage |
| Estimated twinning fraction | No twinning to report. | Xtriage |
| $F_o$,$F_c$ correlation | 0.96 | EDS |
| Total number of atoms | 3756 | wwPDB-VP |
| Average B, all atoms (Å$^2$) | 33.0 | wwPDB-VP |

Xtriage's analysis on translational NCS is as follows: *The largest off-origin peak in the Patterson function is 5.84% of the height of the origin peak. No significant pseudotranslation is detected.*

[1]Intensities estimated from amplitudes.

[2]Theoretical values of $< |L| >$, $< L^2 >$ for acentric reflections are 0.5, 0.333 respectively for untwinned datasets, and 0.375, 0.2 for perfectly twinned datasets.

PRELIMINARY VALIDATION REPORT

### 5 Model quality (i)

#### 5.1 Standard geometry (i)

Bond lengths and bond angles in the following residue types are not validated in this section: ACE

The Z score for a bond length (or angle) is the number of standard deviations the observed value is removed from the expected value. A bond length (or angle) with $|Z| > 5$ is considered an outlier worth inspection. RMSZ is the root-mean-square of all Z scores of the bond lengths (or angles).

| Mol | Chain | Bond lengths | | Bond angles | |
|---|---|---|---|---|---|
| | | RMSZ | $\#\|Z\| > 5$ | RMSZ | $\#\|Z\| > 5$ |
| 1 | A | 0.34 | 0/1084 | 0.54 | 0/1457 |
| 1 | B | 0.36 | 0/707 | 0.53 | 0/951 |
| 2 | D | 0.35 | 0/1104 | 0.55 | 0/1484 |
| 3 | E | 0.36 | 0/694 | 0.55 | 0/937 |
| 4 | F | 0.28 | 0/32 | 0.55 | 0/43 |
| 4 | G | 0.29 | 0/32 | 0.65 | 0/43 |
| All | All | 0.35 | 0/3653 | 0.54 | 0/4915 |

There are no bond length outliers.

There are no bond angle outliers.

There are no chirality outliers.

There are no planarity outliers.

#### 5.2 Too-close contacts (i)

In the following table, the Non-H and H(model) columns list the number of non-hydrogen atoms and hydrogen atoms in the chain respectively. The H(added) column lists the number of hydrogen atoms added and optimized by MolProbity. The Clashes column lists the number of clashes within the asymmetric unit, whereas Symm-Clashes lists symmetry related clashes.

| Mol | Chain | Non-H | H(model) | H(added) | Clashes | Symm-Clashes |
|---|---|---|---|---|---|---|
| 1 | A | 1063 | 0 | 1007 | 5 | 0 |
| 1 | B | 686 | 0 | 670 | 6 | 0 |
| 2 | D | 1085 | 0 | 1016 | 5 | 0 |
| 3 | E | 673 | 0 | 655 | 8 | 0 |
| 4 | F | 35 | 0 | 26 | 2 | 0 |
| 4 | G | 35 | 0 | 26 | 0 | 0 |
| 5 | C | 179 | 0 | 0 | 1 | 0 |
| All | All | 3756 | 0 | 3400 | 20 | 0 |

PRELIMINARY VALIDATION REPORT

The all-atom clashscore is defined as the number of clashes found per 1000 atoms (including hydrogen atoms). The all-atom clashscore for this structure is 3.

All (20) close contacts within the same asymmetric unit are listed below, sorted by their clash magnitude.

| Atom-1 | Atom-2 | Interatomic distance (Å) | Clash overlap (Å) |
|---|---|---|---|
| 1:B:231:SER:HB3 | 1:B:255:LYS:HD2 | 1.79 | 0.64 |
| 2:D:39:MET:O | 2:D:44[A]:ARG:NH1 | 2.32 | 0.63 |
| 1:A:34:ASP:N | 5:C:42:HOH:O | 2.33 | 0.61 |
| 3:E:261:GLU:HG2 | 4:F:2:ASP:OD2 | 2.01 | 0.61 |
| 1:A:100[A]:LEU:HD12 | 1:A:139:LEU:HD22 | 1.85 | 0.58 |
| 3:E:220:ARG:HA | 3:E:226:SER:HA | 1.87 | 0.56 |
| 1:A:48:LEU:HD13 | 1:A:103:VAL:HG21 | 1.88 | 0.56 |
| 1:B:281[B]:MET:HE2 | 3:E:276:PRO:O | 2.08 | 0.53 |
| 2:D:157:LEU:HD22 | 3:E:207:LEU:HD23 | 1.92 | 0.51 |
| 1:B:220:ARG:HA | 1:B:226:SER:HA | 1.92 | 0.50 |
| 3:E:259:ARG:O | 3:E:273:LYS:NZ | 2.27 | 0.49 |
| 1:B:281[B]:MET:HE3 | 3:E:277:CYS:HB2 | 1.94 | 0.49 |
| 2:D:33:LEU:HD12 | 2:D:34:ASP:H | 1.77 | 0.49 |
| 2:D:100:LEU:HD12 | 2:D:139:LEU:HD22 | 1.97 | 0.46 |
| 1:A:100[B]:LEU:HD22 | 1:A:139:LEU:HB3 | 1.98 | 0.45 |
| 3:E:260:SER:OG | 3:E:261:GLU:N | 2.51 | 0.44 |
| 1:B:272:LYS:HD3 | 1:B:272:LYS:HA | 1.79 | 0.44 |
| 1:A:71:ALA:O | 1:A:75:GLU:HG3 | 2.18 | 0.43 |
| 2:D:163:CYS:SG | 4:F:5:ASP:C | 2.98 | 0.41 |
| 1:B:281[B]:MET:CE | 3:E:277:CYS:HB2 | 2.50 | 0.41 |

There are no symmetry-related clashes.

#### 5.3 Torsion angles (i)

##### 5.3.1 Protein backbone (i)

In the following table, the Percentiles column shows the percent Ramachandran outliers of the chain as a percentile score with respect to all X-ray entries followed by that with respect to entries of similar resolution.

The Analysed column shows the number of residues for which the backbone conformation was analysed, and the total number of residues.

| Mol | Chain | Analysed | Favoured | Allowed | Outliers | Percentiles | |
|---|---|---|---|---|---|---|---|
| 1 | A | 140/245 (57%) | 138 (99%) | 2 (1%) | 0 | 100 | 100 |

*Continued on next page...*

PRELIMINARY VALIDATION REPORT

*Continued from previous page...*

| Mol | Chain | Analysed | Favoured | Allowed | Outliers | Percentiles | |
|---|---|---|---|---|---|---|---|
| 1 | B | 83/245 (34%) | 81 (98%) | 2 (2%) | 0 | 100 | 100 |
| 2 | D | 142/142 (100%) | 138 (97%) | 4 (3%) | 0 | 100 | 100 |
| 3 | E | 83/85 (98%) | 82 (99%) | 1 (1%) | 0 | 100 | 100 |
| 4 | F | 3/5 (60%) | 3 (100%) | 0 | 0 | 100 | 100 |
| 4 | G | 3/5 (60%) | 3 (100%) | 0 | 0 | 100 | 100 |
| All | All | 454/727 (62%) | 445 (98%) | 9 (2%) | 0 | 100 | 100 |

There are no Ramachandran outliers to report.

##### 5.3.2 Protein sidechains (i)

In the following table, the Percentiles column shows the percent sidechain outliers of the chain as a percentile score with respect to all X-ray entries followed by that with respect to entries of similar resolution.

The Analysed column shows the number of residues for which the sidechain conformation was analysed, and the total number of residues.

| Mol | Chain | Analysed | Rotameric | Outliers | Percentiles | |
|---|---|---|---|---|---|---|
| 1 | A | 111/217 (51%) | 111 (100%) | 0 | 100 | 100 |
| 1 | B | 74/217 (34%) | 74 (100%) | 0 | 100 | 100 |
| 2 | D | 111/127 (87%) | 108 (97%) | 3 (3%) | 48 | 30 |
| 3 | E | 71/76 (93%) | 70 (99%) | 1 (1%) | 69 | 58 |
| 4 | F | 4/4 (100%) | 4 (100%) | 0 | 100 | 100 |
| 4 | G | 4/4 (100%) | 4 (100%) | 0 | 100 | 100 |
| All | All | 375/645 (58%) | 371 (99%) | 4 (1%) | 83 | 67 |

All (4) residues with a non-rotameric sidechain are listed below:

| Mol | Chain | Res | Type |
|---|---|---|---|
| 2 | D | 44[A] | ARG |
| 2 | D | 44[B] | ARG |
| 2 | D | 80 | SER |
| 3 | E | 261 | GLU |

Some sidechains can be flipped to improve hydrogen bonding and reduce clashes. All (2) such sidechains are listed below:

PRELIMINARY VALIDATION REPORT

| Mol | Chain | Res | Type |
|---|---|---|---|
| 1 | A | 149 | GLN |
| 1 | B | 230 | GLN |

##### 5.3.3 RNA (i)

There are no RNA molecules in this entry.

#### 5.4 Non-standard residues in protein, DNA, RNA chains (i)

There are no non-standard protein/DNA/RNA residues in this entry.

#### 5.5 Carbohydrates (i)

There are no carbohydrates in this entry.

#### 5.6 Ligand geometry (i)

There are no ligands in this entry.

#### 5.7 Other polymers (i)

There are no such residues in this entry.

#### 5.8 Polymer linkage issues (i)

The following chains have linkage breaks:

| Mol | Chain | Number of breaks |
|---|---|---|
| 3 | E | 1 |

All chain breaks are listed below:

| Model | Chain | Residue-1 | Atom-1 | Residue-2 | Atom-2 | Distance (Å) |
|---|---|---|---|---|---|---|
| 1 | E | 261:GLU | C | 270:ASN | N | 5.02 |

PRELIMINARY VALIDATION REPORT

### 6 Fit of model and data (i)

#### 6.1 Protein, DNA and RNA chains (i)

In the following table, the column labelled '#RSRZ> 2' contains the number (and percentage) of RSRZ outliers, followed by percent RSRZ outliers for the chain as percentile scores relative to all X-ray entries and entries of similar resolution. The OWAB column contains the minimum, median, $95^{th}$ percentile and maximum values of the occupancy-weighted average B-factor per residue. The column labelled 'Q< 0.9' lists the number of (and percentage) of residues with an average occupancy less than 0.9.

| Mol | Chain | Analysed | <RSRZ> | #RSRZ>2 | | | OWAB(Å$^2$) | Q<0.9 |
|---|---|---|---|---|---|---|---|---|
| 1 | A | 139/245 (56%) | -0.16 | 2 (1%) | 75 | 75 | 22, 31, 48, 58 | 0 |
| 1 | B | 87/245 (35%) | 0.00 | 0 | 100 | 100 | 22, 31, 49, 66 | 0 |
| 2 | D | 142/142 (100%) | 0.05 | 4 (2%) | 53 | 50 | 23, 31, 48, 65 | 0 |
| 3 | E | 85/85 (100%) | 0.28 | 3 (3%) | 44 | 40 | 23, 31, 51, 65 | 0 |
| 4 | F | 4/5 (80%) | 0.09 | 0 | 100 | 100 | 34, 38, 43, 44 | 0 |
| 4 | G | 4/5 (80%) | -0.01 | 0 | 100 | 100 | 30, 33, 39, 42 | 0 |
| All | All | 461/727 (63%) | 0.02 | 9 (1%) | 65 | 64 | 22, 31, 49, 66 | 0 |

All (9) RSRZ outliers are listed below:

| Mol | Chain | Res | Type | RSRZ |
|---|---|---|---|---|
| 1 | A | 172 | VAL | 5.6 |
| 3 | E | 261 | GLU | 2.9 |
| 2 | D | 172 | VAL | 2.8 |
| 3 | E | 270 | ASN | 2.5 |
| 2 | D | 33 | LEU | 2.4 |
| 2 | D | 34 | ASP | 2.3 |
| 3 | E | 259 | ARG | 2.1 |
| 1 | A | 171 | GLY | 2.1 |
| 2 | D | 136 | ILE | 2.1 |

#### 6.2 Non-standard residues in protein, DNA, RNA chains (i)

There are no non-standard protein/DNA/RNA residues in this entry.

#### 6.3 Carbohydrates (i)

There are no carbohydrates in this entry.

PRELIMINARY VALIDATION REPORT

#### 6.4 Ligands (i)

There are no ligands in this entry.

#### 6.5 Other polymers (i)

There are no such residues in this entry.

PRELIMINARY VALIDATION REPORT

### Preliminary Full wwPDB X-ray Structure Validation Report (i)

Jul 8, 2019 – 01:02 PM EDT

| | | |
|---|---|---|
| Deposition ID | : | D_1000242855 |
| PDB ID | : | *(not yet assigned)* |

This is a Preliminary Full wwPDB X-ray Structure Validation Report.

This report is produced by the wwPDB Deposition System during initial deposition but before annotation of the structure.

We welcome your comments at **
A user guide is available at
https://www.wwpdb.org/validation/2017/XrayValidationReportHelp
with specific help available everywhere you see the (i) symbol.

The following versions of software and data (see references (i)) were used in the production of this report:

| | | |
|---|---|---|
| MolProbity | : | 4.02b-467 |
| Xtriage (Phenix) | : | 1.13 |
| EDS | : | 2.4 |
| Percentile statistics | : | 20171227.v01 (using entries in the PDB archive December 27th 2017) |
| Refmac | : | 5.8.0158 |
| CCP4 | : | 7.0 (Gargrove) |
| Ideal geometry (proteins) | : | Engh & Huber (2001) |
| Ideal geometry (DNA, RNA) | : | Parkinson et al. (1996) |
| Validation Pipeline (wwPDB-VP) | : | 2.4 |

PRELIMINARY VALIDATION REPORT

### 1 Overall quality at a glance (i)

The following experimental techniques were used to determine the structure:
*X-RAY DIFFRACTION*

The reported resolution of this entry is 2.61 Å.

Percentile scores (ranging between 0-100) for global validation metrics of the entry are shown in the following graphic. The table shows the number of entries on which the scores are based.

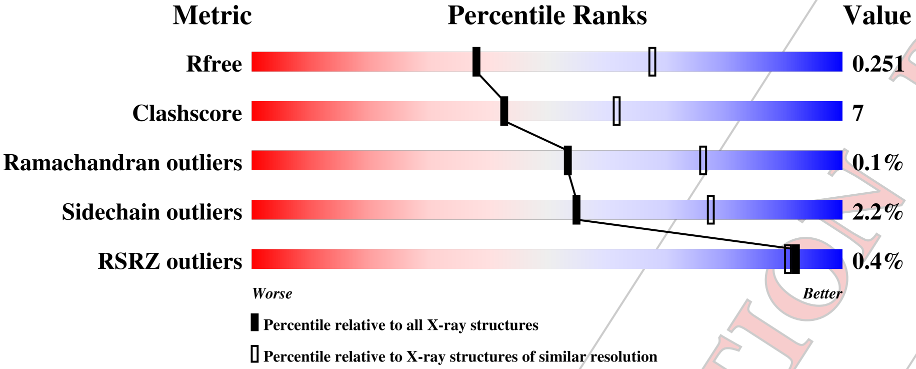


| Metric | Whole archive (#Entries) | Similar resolution (#Entries, resolution range(Å)) |
|---|---|---|
| $R_{free}$ | 111664 | 3285 (2.64-2.60) |
| Clashscore | 122126 | 3641 (2.64-2.60) |
| Ramachandran outliers | 120053 | 3586 (2.64-2.60) |
| Sidechain outliers | 120020 | 3586 (2.64-2.60) |
| RSRZ outliers | 108989 | 3218 (2.64-2.60) |

The table below summarises the geometric issues observed across the polymeric chains and their fit to the electron density. The red, orange, yellow and green segments on the lower bar indicate the fraction of residues that contain outliers for >=3, 2, 1 and 0 types of geometric quality criteria. A grey segment represents the fraction of residues that are not modelled. The numeric value for each fraction is indicated below the corresponding segment, with a dot representing fractions <=5% The upper red bar (where present) indicates the fraction of residues that have poor fit to the electron density. The numeric value is given above the bar.

| Mol | Chain | Length | Quality of chain |
|---|---|---|---|
| 1 | A | 151 | 77% 23% • |
| 1 | G | 151 | 89% 11% |
| 2 | B | 94 | % 87% 13% |
| 2 | H | 94 | 88% 11% • |
| 3 | E | 4 | 75% 25% |

*Continued on next page...*

WORLDWIDE PDB PROTEIN DATA BANK

PRELIMINARY VALIDATION REPORT

*Continued from previous page...*

| Mol | Chain | Length | Quality of chain |
|---|---|---|---|
| 3 | I | 4 | 75% 25% |
| 3 | L | 4 | 25% 75% |
| 4 | C | 145 | 82% 18% |
| 5 | D | 95 | % 77% 21% •• |
| 6 | J | 144 | % 76% 24% |
| 7 | K | 96 | % 74% 18% 8% |

PRELIMINARY VALIDATION REPORT

### 2 Entry composition (i)

There are 9 unique types of molecules in this entry. The entry contains 7697 atoms, of which 0 are hydrogens and 0 are deuteriums.

In the tables below, the ZeroOcc column contains the number of atoms modelled with zero occupancy, the AltConf column contains the number of residues with at least one atom in alternate conformation and the Trace column contains the number of residues modelled with at most 2 atoms.

- Molecule 1 is a protein called Ancestral Caspase-6 Large Subunit.

| Mol | Chain | Residues | Atoms | | | | | ZeroOcc | AltConf | Trace |
|---|---|---|---|---|---|---|---|---|---|---|
| 1 | A | 151 | Total 1185 | C 746 | N 206 | O 226 | S 7 | 0 | 0 | 0 |
| 1 | G | 151 | Total 1183 | C 749 | N 205 | O 223 | S 6 | 0 | 1 | 0 |

- Molecule 2 is a protein called Ancestral Caspase-6 small subunit.

| Mol | Chain | Residues | Atoms | | | | | ZeroOcc | AltConf | Trace |
|---|---|---|---|---|---|---|---|---|---|---|
| 2 | B | 94 | Total 751 | C 488 | N 118 | O 136 | S 9 | 5 | 2 | 0 |
| 2 | H | 94 | Total 749 | C 487 | N 118 | O 136 | S 8 | 0 | 0 | 0 |

- Molecule 3 is a protein called VAL-GLU-ILE-ASP Inhibitor.

| Mol | Chain | Residues | Atoms | | | | ZeroOcc | AltConf | Trace |
|---|---|---|---|---|---|---|---|---|---|
| 3 | E | 4 | Total 32 | C 20 | N 4 | O 8 | 0 | 0 | 0 |
| 3 | I | 4 | Total 32 | C 20 | N 4 | O 8 | 0 | 0 | 0 |
| 3 | L | 4 | Total 32 | C 20 | N 4 | O 8 | 0 | 0 | 0 |

- Molecule 4 is a protein called Ancestral Caspase-6 Large Subunit.

| Mol | Chain | Residues | Atoms | | | | | ZeroOcc | AltConf | Trace |
|---|---|---|---|---|---|---|---|---|---|---|
| 4 | C | 145 | Total 1107 | C 695 | N 190 | O 216 | S 6 | 0 | 0 | 0 |

- Molecule 5 is a protein called Ancestral Caspase-6 Small Subunit.

PRELIMINARY VALIDATION REPORT

| Mol | Chain | Residues | Atoms | | | | | ZeroOcc | AltConf | Trace |
|---|---|---|---|---|---|---|---|---|---|---|
| | | | Total | C | N | O | S | | | |
| 5 | D | 94 | 739 | 483 | 114 | 134 | 8 | 0 | 0 | 0 |

- Molecule 6 is a protein called Ancestral Caspase-6 Large Subunit.

| Mol | Chain | Residues | Atoms | | | | | ZeroOcc | AltConf | Trace |
|---|---|---|---|---|---|---|---|---|---|---|
| | | | Total | C | N | O | S | | | |
| 6 | J | 144 | 1089 | 687 | 193 | 203 | 6 | 5 | 1 | 0 |

- Molecule 7 is a protein called Ancestral Caspase-6 Small Subunit.

| Mol | Chain | Residues | Atoms | | | | | ZeroOcc | AltConf | Trace |
|---|---|---|---|---|---|---|---|---|---|---|
| | | | Total | C | N | O | S | | | |
| 7 | K | 88 | 691 | 451 | 110 | 123 | 7 | 0 | 0 | 0 |

- Molecule 8 is VALINE (three-letter code: VAL) (formula: $C_5H_{11}NO_2$).

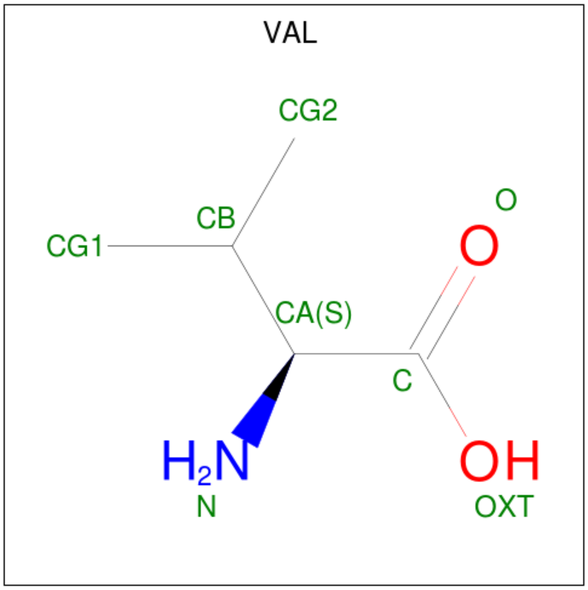


| Mol | Chain | Residues | Atoms | | | | ZeroOcc | AltConf |
|---|---|---|---|---|---|---|---|---|
| | | | Total | C | N | O | | |
| 8 | F | 1 | 7 | 5 | 1 | 1 | 0 | 0 |

- Molecule 9 is water.

| Mol | Chain | Residues | Atoms | | ZeroOcc | AltConf |
|---|---|---|---|---|---|---|
| | | | Total | O | | |
| 9 | M | 100 | 100 | 100 | 0 | 0 |

PRELIMINARY VALIDATION REPORT

### 3 Residue-property plots (i)

These plots are drawn for all protein, RNA and DNA chains in the entry. The first graphic for a chain summarises the proportions of the various outlier classes displayed in the second graphic. The second graphic shows the sequence view annotated by issues in geometryand electron density. Residues are color-coded according to the number of geometric quality criteria for which they contain at least one outlier: green = 0, yellow = 1, orange = 2 and red = 3 or more. A red dot above a residue indicates a poor fit to the electron density (RSRZ > 2). Stretches of 2 or more consecutive residues without any outlier are shown as a green connector. Residues present in the sample, but not in the model, are shown in grey.

- Molecule 1: Ancestral Caspase-6 Large Subunit

Chain A:

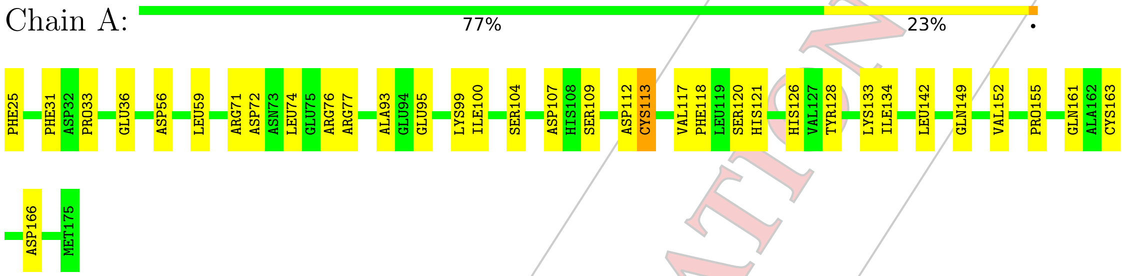


- Molecule 1: Ancestral Caspase-6 Large Subunit

Chain G:

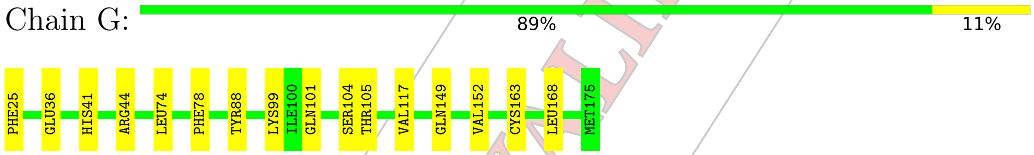


- Molecule 2: Ancestral Caspase-6 small subunit

Chain B:

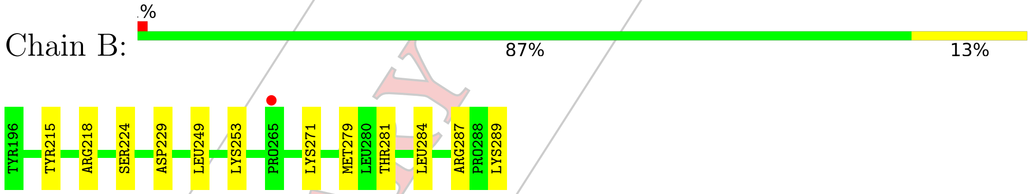


- Molecule 2: Ancestral Caspase-6 small subunit

Chain H:

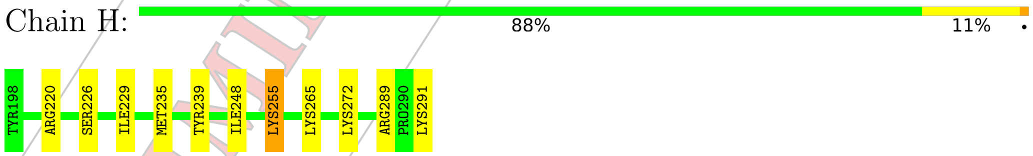


- Molecule 3: VAL-GLU-ILE-ASP Inhibitor

Chain E:

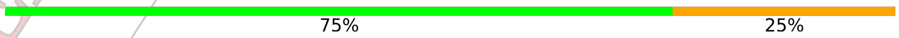


PRELIMINARY VALIDATION REPORT

VAL301 ASP304

• Molecule 3: VAL-GLU-ILE-ASP Inhibitor

Chain I: 75% 25%

VAL301 ASP304

• Molecule 3: VAL-GLU-ILE-ASP Inhibitor

Chain L: 25% 75%

VAL301 GLU302 ILE303 ASP304

• Molecule 4: Ancestral Caspase-6 Large Subunit

Chain C: 82% 18%

PHE31 ASP32 PRO33 GLU36 ARG44 PHE55 ARG64 LEU74 GLU75 ARG76 ARG77 PHE78 LEU81 VAL85 SER92 GLU95 ILE100 SER104 VAL117 PHE118 LEU119 HIS126 LYS133 LEU151 LYS154 PRO155 LYS156 GLN161 LEU168 ASP169 MET175

• Molecule 5: Ancestral Caspase-6 Small Subunit

Chain D: % 77% 21% ••

VAL197 TYR198 PRO201 TYR217 SER218 TYR219 ARG GLU221 THR222 VAL223 ASN224 GLY225 SER226 GLN230 GLU234 TYR239 LEU243 GLU244 PHE245 THR246 GLU247 LEU251 ARG254 LYS255 LEU258 LYS272 ARG289 PRO290 LYS291

• Molecule 6: Ancestral Caspase-6 Large Subunit

Chain J: % 76% 24%

ASP32 ARG43 ARG44 ASP56 TRP57 LYS58 LEU59 GLY60 LEU61 ARG64 THR67 ASP70 LEU74 ARG77 PHE78 GLN79 GLY82 ALA93 GLU98 LYS99 ILE100 HIS108 SER109 ASP110 ALA111 ASP112 LEU119 SER120 HIS121 HIS126 ASP131 ALA132 LYS133 ILE134 GLU135 GLU138 LEU142 ASP146 GLN149 SER150 LEU151 VAL152

LEU168 MET175

• Molecule 7: Ancestral Caspase-6 Small Subunit

Chain K: % 74% 18% 8%

PRELIMINARY VALIDATION REPORT

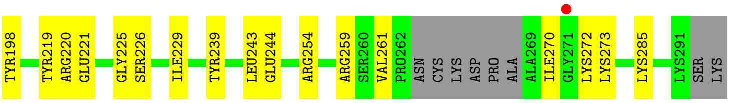


PRELIMINARY VALIDATION REPORT

### 4 Data and refinement statistics (i)

| Property | Value | Source |
|---|---|---|
| Space group | P 21 21 21 | Depositor |
| Cell constants<br>a, b, c, $\alpha$, $\beta$, $\gamma$ | 84.31Å 88.54Å 141.03Å<br>90.00° 90.00° 90.00° | Depositor |
| Resolution (Å) | 42.24 – 2.61<br>42.24 – 2.61 | Depositor<br>EDS |
| % Data completeness<br>(in resolution range) | 99.7 (42.24-2.61)<br>99.7 (42.24-2.61) | Depositor<br>EDS |
| $R_{merge}$ | (Not available) | Depositor |
| $R_{sym}$ | (Not available) | Depositor |
| $< I/\sigma(I) >$ [1] | 2.11 (at 2.61Å) | Xtriage |
| Refinement program | phenix.refine 1.15.2_3472, PHENIX 1.15.2_3472 | Depositor |
| R, $R_{free}$ | 0.177 , 0.251<br>0.177 , 0.251 | Depositor<br>DCC |
| $R_{free}$ test set | 1999 reflections (6.11%) | wwPDB-VP |
| Wilson B-factor (Å$^2$) | 51.8 | Xtriage |
| Anisotropy | 0.639 | Xtriage |
| Bulk solvent $k_{sol}$(e/Å$^3$), $B_{sol}$(Å$^2$) | 0.31 , 46.3 | EDS |
| L-test for twinning[2] | $< \|L\| > = 0.49$, $< L^2 > = 0.33$ | Xtriage |
| Estimated twinning fraction | 0.023 for k,h,-l | Xtriage |
| $F_o$,$F_c$ correlation | 0.95 | EDS |
| Total number of atoms | 7697 | wwPDB-VP |
| Average B, all atoms (Å$^2$) | 54.0 | wwPDB-VP |

Xtriage's analysis on translational NCS is as follows: *The largest off-origin peak in the Patterson function is 4.69% of the height of the origin peak. No significant pseudotranslation is detected.*

[1] Intensities estimated from amplitudes.
[2] Theoretical values of $< |L| >$, $< L^2 >$ for acentric reflections are 0.5, 0.333 respectively for untwinned datasets, and 0.375, 0.2 for perfectly twinned datasets.

PRELIMINARY VALIDATION REPORT

### 5 Model quality (i)

#### 5.1 Standard geometry (i)

The Z score for a bond length (or angle) is the number of standard deviations the observed value is removed from the expected value. A bond length (or angle) with $|Z| > 5$ is considered an outlier worth inspection. RMSZ is the root-mean-square of all Z scores of the bond lengths (or angles).

| Mol | Chain | Bond lengths | | Bond angles | |
|---|---|---|---|---|---|
| | | RMSZ | #\|Z\| >5 | RMSZ | #\|Z\| >5 |
| 1 | A | 0.45 | 1/1207 (0.1%) | 0.57 | 0/1626 |
| 1 | G | 0.41 | 0/1208 | 0.56 | 0/1628 |
| 2 | B | 0.43 | 0/775 | 0.62 | 0/1048 |
| 2 | H | 0.48 | 0/767 | 0.60 | 0/1034 |
| 3 | E | 0.34 | 0/31 | 0.62 | 0/41 |
| 3 | I | 0.62 | 0/31 | 0.82 | 0/41 |
| 3 | L | 0.39 | 0/31 | 0.46 | 0/41 |
| 4 | C | 0.40 | 0/1127 | 0.60 | 0/1525 |
| 5 | D | 0.48 | 0/756 | 0.61 | 0/1020 |
| 6 | J | 0.40 | 0/1111 | 0.56 | 0/1503 |
| 7 | K | 0.40 | 0/707 | 0.53 | 0/955 |
| All | All | 0.43 | 1/7751 (0.0%) | 0.58 | 0/10462 |

All (1) bond length outliers are listed below:

| Mol | Chain | Res | Type | Atoms | Z | Observed(Å) | Ideal(Å) |
|---|---|---|---|---|---|---|---|
| 1 | A | 113 | CYS | CB-SG | 5.06 | 1.90 | 1.82 |

There are no bond angle outliers.

There are no chirality outliers.

There are no planarity outliers.

#### 5.2 Too-close contacts (i)

In the following table, the Non-H and H(model) columns list the number of non-hydrogen atoms and hydrogen atoms in the chain respectively. The H(added) column lists the number of hydrogen atoms added and optimized by MolProbity. The Clashes column lists the number of clashes within the asymmetric unit, whereas Symm-Clashes lists symmetry related clashes.

| Mol | Chain | Non-H | H(model) | H(added) | Clashes | Symm-Clashes |
|---|---|---|---|---|---|---|
| 1 | A | 1185 | 0 | 1111 | 26 | 0 |

*Continued on next page...*

PRELIMINARY VALIDATION REPORT

*Continued from previous page...*

| Mol | Chain | Non-H | H(model) | H(added) | Clashes | Symm-Clashes |
|---|---|---|---|---|---|---|
| 1 | G | 1183 | 0 | 1113 | 12 | 0 |
| 2 | B | 751 | 0 | 745 | 12 | 0 |
| 2 | H | 749 | 0 | 751 | 11 | 0 |
| 3 | E | 32 | 0 | 29 | 4 | 0 |
| 3 | I | 32 | 0 | 29 | 3 | 0 |
| 3 | L | 32 | 0 | 29 | 4 | 0 |
| 4 | C | 1107 | 0 | 1013 | 18 | 0 |
| 5 | D | 739 | 0 | 732 | 14 | 0 |
| 6 | J | 1089 | 0 | 1017 | 24 | 0 |
| 7 | K | 691 | 0 | 678 | 12 | 0 |
| 8 | F | 7 | 0 | 8 | 0 | 0 |
| 9 | M | 100 | 0 | 0 | 2 | 0 |
| All | All | 7697 | 0 | 7255 | 110 | 0 |

The all-atom clashscore is defined as the number of clashes found per 1000 atoms (including hydrogen atoms). The all-atom clashscore for this structure is 7.

All (110) close contacts within the same asymmetric unit are listed below, sorted by their clash magnitude.

| Atom-1 | Atom-2 | Interatomic distance (Å) | Clash overlap (Å) |
|---|---|---|---|
| 7:K:261:VAL:HG12 | 7:K:272:LYS:H | 1.39 | 0.87 |
| 2:H:220:ARG:HH21 | 3:I:304:ASP:HB3 | 1.43 | 0.82 |
| 1:A:107:ASP:OD1 | 1:A:109:SER:OG | 2.06 | 0.73 |
| 2:B:218:ARG:HH21 | 3:E:304:ASP:HB3 | 1.58 | 0.68 |
| 1:A:36:GLU:OE1 | 2:B:287:ARG:NH2 | 2.26 | 0.68 |
| 2:H:220:ARG:HA | 2:H:226:SER:HA | 1.75 | 0.68 |
| 7:K:220:ARG:HA | 7:K:226:SER:HA | 1.77 | 0.67 |
| 1:A:100:ILE:HG22 | 1:A:142:LEU:HD12 | 1.79 | 0.65 |
| 4:C:168:LEU:HD13 | 5:D:217:TYR:HB2 | 1.78 | 0.64 |
| 7:K:259:ARG:O | 7:K:273:LYS:NZ | 2.24 | 0.63 |
| 4:C:126:HIS:HB3 | 4:C:133:LYS:HG2 | 1.81 | 0.62 |
| 1:G:101:GLN:O | 1:G:105:THR:HG22 | 2.00 | 0.62 |
| 5:D:244:GLU:OE1 | 5:D:246:THR:OG1 | 2.16 | 0.62 |
| 2:B:218:ARG:HA | 2:B:224:SER:HA | 1.80 | 0.62 |
| 7:K:220:ARG:O | 3:L:302:GLU:N | 2.27 | 0.60 |
| 4:C:44:ARG:HG3 | 4:C:44:ARG:HH11 | 1.66 | 0.60 |
| 1:A:104:SER:HB2 | 1:A:142:LEU:HB3 | 1.84 | 0.60 |
| 1:G:163:CYS:SG | 3:I:304:ASP:C | 2.80 | 0.59 |
| 4:C:151:LEU:HA | 4:C:154:LYS:HD2 | 1.83 | 0.59 |
| 1:A:33:PRO:HA | 5:D:251:LEU:HD21 | 1.84 | 0.59 |
| 2:H:239:TYR:HB2 | 2:H:248:ILE:HD11 | 1.85 | 0.58 |

*Continued on next page...*

PRELIMINARY VALIDATION REPORT

*Continued from previous page...*

| Atom-1 | Atom-2 | Interatomic distance (Å) | Clash overlap (Å) |
|---|---|---|---|
| 6:J:43:ARG:NH1 | 6:J:110:ASP:HB2 | 2.18 | 0.58 |
| 2:H:220:ARG:HH21 | 3:I:304:ASP:CB | 2.16 | 0.58 |
| 4:C:36:GLU:OE1 | 5:D:289:ARG:NH2 | 2.37 | 0.58 |
| 6:J:168:LEU:HD23 | 7:K:219:TYR:OH | 2.04 | 0.58 |
| 4:C:92:SER:HB3 | 4:C:95:GLU:HG3 | 1.86 | 0.57 |
| 1:G:41:HIS:O | 2:H:291:LYS:NZ | 2.38 | 0.56 |
| 1:A:74:LEU:HD13 | 1:A:117:VAL:HG11 | 1.89 | 0.55 |
| 7:K:261:VAL:HG13 | 7:K:270:ILE:HA | 1.88 | 0.55 |
| 4:C:119:LEU:HD23 | 4:C:161:GLN:HB3 | 1.88 | 0.54 |
| 4:C:75:GLU:HG3 | 4:C:85:VAL:HG11 | 1.89 | 0.54 |
| 1:G:168:LEU:HD22 | 2:H:272:LYS:HG3 | 1.90 | 0.54 |
| 1:A:161:GLN:NE2 | 9:M:14:HOH:O | 2.30 | 0.54 |
| 7:K:221:GLU:HA | 3:L:301:VAL:HA | 1.90 | 0.54 |
| 1:A:166:ASP:HA | 2:B:215:TYR:CE1 | 2.42 | 0.54 |
| 4:C:74:LEU:HD13 | 4:C:117:VAL:HG11 | 1.90 | 0.53 |
| 1:A:31:PHE:HE2 | 5:D:258:LEU:HD12 | 1.72 | 0.53 |
| 6:J:77:ARG:NE | 6:J:77:ARG:HA | 2.23 | 0.53 |
| 1:A:155:PRO:HG2 | 2:B:284:LEU:HD13 | 1.89 | 0.53 |
| 1:G:88:TYR:CZ | 1:G:99:LYS:HD3 | 2.45 | 0.52 |
| 1:A:128:TYR:CE1 | 1:A:133:LYS:HG2 | 2.45 | 0.52 |
| 2:B:281:THR:HB | 5:D:254:ARG:HG3 | 1.92 | 0.51 |
| 1:G:44:ARG:HG3 | 1:G:44:ARG:HH11 | 1.75 | 0.51 |
| 6:J:44:ARG:O | 6:J:112:ASP:N | 2.41 | 0.51 |
| 4:C:44:ARG:NE | 4:C:81:LEU:O | 2.34 | 0.51 |
| 1:G:74:LEU:HD13 | 1:G:117:VAL:HG11 | 1.93 | 0.51 |
| 5:D:234:GLU:OE1 | 5:D:255:LYS:HE3 | 2.10 | 0.51 |
| 5:D:244:GLU:HB3 | 5:D:247:GLU:HG3 | 1.91 | 0.51 |
| 6:J:108:HIS:H | 6:J:150:SER:HB2 | 1.75 | 0.51 |
| 2:B:249:LEU:HD21 | 4:C:33:PRO:HA | 1.93 | 0.50 |
| 1:G:149:GLN:HA | 1:G:152:VAL:HG23 | 1.93 | 0.50 |
| 6:J:149:GLN:HA | 6:J:152:VAL:HG23 | 1.93 | 0.50 |
| 6:J:93:ALA:HB1 | 6:J:134:ILE:HD11 | 1.94 | 0.50 |
| 1:A:112:ASP:OD2 | 2:B:289:LYS:NZ | 2.38 | 0.50 |
| 1:A:76:ARG:HG2 | 1:A:77:ARG:NH1 | 2.26 | 0.49 |
| 1:A:163:CYS:SG | 3:E:304:ASP:C | 2.91 | 0.49 |
| 4:C:55:PHE:HE2 | 4:C:64:ARG:HG3 | 1.77 | 0.49 |
| 6:J:56:ASP:OD1 | 6:J:58:LYS:HE3 | 2.13 | 0.48 |
| 1:A:104:SER:CB | 1:A:142:LEU:HB3 | 2.44 | 0.48 |
| 2:B:229:ASP:CG | 2:B:253:LYS:HG2 | 2.34 | 0.48 |
| 6:J:64:ARG:HE | 3:L:304:ASP:HB2 | 1.78 | 0.48 |
| 7:K:220:ARG:HH21 | 3:L:304:ASP:HB3 | 1.78 | 0.48 |

*Continued on next page...*

PRELIMINARY VALIDATION REPORT

*Continued from previous page...*

| Atom-1 | Atom-2 | Interatomic distance (Å) | Clash overlap (Å) |
|---|---|---|---|
| 6:J:100:ILE:HG22 | 6:J:142:LEU:HD12 | 1.96 | 0.48 |
| 1:A:126:HIS:CE1 | 1:A:133:LYS:HD2 | 2.49 | 0.47 |
| 4:C:154:LYS:O | 4:C:156:LYS:HE3 | 2.15 | 0.46 |
| 1:A:71:ARG:NH2 | 1:A:72:ASP:OD1 | 2.42 | 0.46 |
| 6:J:64:ARG:NH1 | 6:J:67:THR:HG21 | 2.30 | 0.46 |
| 1:A:36:GLU:CD | 2:B:287:ARG:HH22 | 2.19 | 0.46 |
| 1:A:120:SER:OG | 1:A:121:HIS:N | 2.49 | 0.46 |
| 4:C:78:PHE:HE2 | 4:C:117:VAL:HG21 | 1.81 | 0.46 |
| 7:K:221:GLU:N | 7:K:225:GLY:O | 2.48 | 0.45 |
| 1:A:113:CYS:HB2 | 1:A:155:PRO:O | 2.17 | 0.45 |
| 6:J:59:LEU:HD21 | 6:J:131:ASP:O | 2.17 | 0.45 |
| 7:K:239:TYR:HB3 | 7:K:243:LEU:HG | 1.97 | 0.45 |
| 1:G:36:GLU:OE1 | 2:H:289:ARG:NH1 | 2.48 | 0.44 |
| 6:J:74:LEU:HD11 | 7:K:229:ILE:HD12 | 1.99 | 0.44 |
| 5:D:222:THR:O | 5:D:224:ASN:N | 2.41 | 0.44 |
| 7:K:244:GLU:HG3 | 7:K:285:LYS:HB3 | 1.98 | 0.44 |
| 4:C:168:LEU:HB3 | 5:D:272:LYS:HD3 | 1.99 | 0.44 |
| 4:C:77:ARG:HA | 4:C:77:ARG:NE | 2.32 | 0.44 |
| 1:G:74:LEU:HD22 | 1:G:78:PHE:HE2 | 1.83 | 0.44 |
| 6:J:108:HIS:H | 6:J:150:SER:CB | 2.31 | 0.44 |
| 6:J:126:HIS:CD2 | 6:J:133:LYS:HE3 | 2.53 | 0.44 |
| 1:A:126:HIS:NE2 | 1:A:133:LYS:HD2 | 2.33 | 0.44 |
| 5:D:221:GLU:O | 5:D:223:VAL:N | 2.52 | 0.43 |
| 2:H:248:ILE:HD13 | 2:H:248:ILE:HA | 1.88 | 0.43 |
| 5:D:226:SER:O | 5:D:230:GLN:HB3 | 2.18 | 0.43 |
| 2:H:235:MET:CE | 2:H:255:LYS:HD2 | 2.49 | 0.43 |
| 6:J:77:ARG:HE | 6:J:77:ARG:HA | 1.83 | 0.43 |
| 6:J:135:GLU:O | 6:J:138:GLU:HB2 | 2.19 | 0.43 |
| 1:A:59:LEU:HD23 | 1:A:59:LEU:HA | 1.78 | 0.42 |
| 2:B:218:ARG:NH2 | 3:E:304:ASP:HB3 | 2.31 | 0.42 |
| 1:A:149:GLN:HA | 1:A:152:VAL:HG23 | 2.01 | 0.41 |
| 6:J:64:ARG:CZ | 6:J:67:THR:HG21 | 2.49 | 0.41 |
| 1:A:95:GLU:O | 1:A:99:LYS:HG2 | 2.21 | 0.41 |
| 4:C:44:ARG:HG3 | 4:C:44:ARG:NH1 | 2.33 | 0.41 |
| 2:H:265:LYS:HE2 | 2:H:265:LYS:HB3 | 1.82 | 0.41 |
| 6:J:67:THR:H | 6:J:67:THR:HG23 | 1.63 | 0.41 |
| 5:D:239:TYR:HB3 | 5:D:243:LEU:HD12 | 2.03 | 0.41 |
| 1:G:44:ARG:HG3 | 1:G:44:ARG:NH1 | 2.36 | 0.41 |
| 2:B:271:LYS:HB3 | 5:D:201:PRO:HG3 | 2.02 | 0.40 |
| 4:C:100:ILE:O | 4:C:104:SER:HB3 | 2.21 | 0.40 |
| 6:J:44:ARG:NH2 | 9:M:143:HOH:O | 2.43 | 0.40 |

*Continued on next page...*

PRELIMINARY VALIDATION REPORT

*Continued from previous page...*

| Atom-1 | Atom-2 | Interatomic distance (Å) | Clash overlap (Å) |
|---|---|---|---|
| 6:J:67:THR:HA | 6:J:70:ASP:HB2 | 2.03 | 0.40 |
| 1:G:74:LEU:HD11 | 2:H:229:ILE:HD12 | 2.04 | 0.40 |
| 6:J:61:LEU:HD13 | 6:J:121:HIS:CD2 | 2.56 | 0.40 |
| 1:A:121:HIS:CE1 | 3:E:304:ASP:HA | 2.56 | 0.40 |
| 6:J:79:GLN:O | 6:J:82:GLY:N | 2.49 | 0.40 |
| 1:A:93:ALA:HA | 1:A:134:ILE:HD11 | 2.03 | 0.40 |
| 6:J:151:LEU:HA | 6:J:151:LEU:HD23 | 1.87 | 0.40 |

There are no symmetry-related clashes.

#### 5.3 Torsion angles (i)

##### 5.3.1 Protein backbone (i)

In the following table, the Percentiles column shows the percent Ramachandran outliers of the chain as a percentile score with respect to all X-ray entries followed by that with respect to entries of similar resolution.

The Analysed column shows the number of residues for which the backbone conformation was analysed, and the total number of residues.

| Mol | Chain | Analysed | Favoured | Allowed | Outliers | Percentiles | |
|---|---|---|---|---|---|---|---|
| 1 | A | 149/151 (99%) | 144 (97%) | 5 (3%) | 0 | 100 | 100 |
| 1 | G | 150/151 (99%) | 143 (95%) | 7 (5%) | 0 | 100 | 100 |
| 2 | B | 94/94 (100%) | 89 (95%) | 5 (5%) | 0 | 100 | 100 |
| 2 | H | 92/94 (98%) | 89 (97%) | 3 (3%) | 0 | 100 | 100 |
| 3 | E | 2/4 (50%) | 2 (100%) | 0 | 0 | 100 | 100 |
| 3 | I | 2/4 (50%) | 2 (100%) | 0 | 0 | 100 | 100 |
| 3 | L | 2/4 (50%) | 2 (100%) | 0 | 0 | 100 | 100 |
| 4 | C | 143/145 (99%) | 136 (95%) | 7 (5%) | 0 | 100 | 100 |
| 5 | D | 90/95 (95%) | 87 (97%) | 2 (2%) | 1 (1%) | 16 | 30 |
| 6 | J | 143/144 (99%) | 132 (92%) | 11 (8%) | 0 | 100 | 100 |
| 7 | K | 84/96 (88%) | 82 (98%) | 2 (2%) | 0 | 100 | 100 |
| All | All | 951/982 (97%) | 908 (96%) | 42 (4%) | 1 (0%) | 53 | 76 |

All (1) Ramachandran outliers are listed below:

PRELIMINARY VALIDATION REPORT

| Mol | Chain | Res | Type |
|---|---|---|---|
| 5 | D | 222 | THR |

##### 5.3.2 Protein sidechains (i)

In the following table, the Percentiles column shows the percent sidechain outliers of the chain as a percentile score with respect to all X-ray entries followed by that with respect to entries of similar resolution.

The Analysed column shows the number of residues for which the sidechain conformation was analysed, and the total number of residues.

| Mol | Chain | Analysed | Rotameric | Outliers | Percentiles | |
|---|---|---|---|---|---|---|
| 1 | A | 120/131 (92%) | 117 (98%) | 3 (2%) | 50 | 75 |
| 1 | G | 118/131 (90%) | 116 (98%) | 2 (2%) | 63 | 82 |
| 2 | B | 81/83 (98%) | 79 (98%) | 2 (2%) | 50 | 75 |
| 2 | H | 81/83 (98%) | 80 (99%) | 1 (1%) | 74 | 88 |
| 3 | E | 4/4 (100%) | 3 (75%) | 1 (25%) | 0 | 1 |
| 3 | I | 4/4 (100%) | 3 (75%) | 1 (25%) | 0 | 1 |
| 3 | L | 4/4 (100%) | 4 (100%) | 0 | 100 | 100 |
| 4 | C | 110/125 (88%) | 108 (98%) | 2 (2%) | 62 | 81 |
| 5 | D | 78/84 (93%) | 77 (99%) | 1 (1%) | 71 | 87 |
| 6 | J | 106/124 (86%) | 102 (96%) | 4 (4%) | 36 | 62 |
| 7 | K | 72/85 (85%) | 70 (97%) | 2 (3%) | 47 | 72 |
| All | All | 778/858 (91%) | 759 (98%) | 19 (2%) | 55 | 76 |

All (19) residues with a non-rotameric sidechain are listed below:

| Mol | Chain | Res | Type |
|---|---|---|---|
| 1 | A | 25 | PHE |
| 1 | A | 56 | ASP |
| 1 | A | 118 | PHE |
| 2 | B | 279[A] | MET |
| 2 | B | 279[B] | MET |
| 3 | E | 304 | ASP |
| 1 | G | 25 | PHE |
| 1 | G | 104 | SER |
| 2 | H | 255 | LYS |
| 3 | I | 304 | ASP |
| 4 | C | 31 | PHE |

*Continued on next page...*

PRELIMINARY VALIDATION REPORT

*Continued from previous page...*

| Mol | Chain | Res | Type |
|---|---|---|---|
| 4 | C | 169 | ASP |
| 5 | D | 198 | TYR |
| 6 | J | 98 | GLU |
| 6 | J | 119[A] | LEU |
| 6 | J | 119[B] | LEU |
| 6 | J | 146 | ASP |
| 7 | K | 198 | TYR |
| 7 | K | 254 | ARG |

Some sidechains can be flipped to improve hydrogen bonding and reduce clashes. There are no such sidechains identified.

###### 5.3.3 RNA (i)

There are no RNA molecules in this entry.

##### 5.4 Non-standard residues in protein, DNA, RNA chains (i)

There are no non-standard protein/DNA/RNA residues in this entry.

##### 5.5 Carbohydrates (i)

There are no carbohydrates in this entry.

##### 5.6 Ligand geometry (i)

1 ligand is modelled in this entry.

There are no bond length outliers.

There are no bond angle outliers.

There are no chirality outliers.

There are no torsion outliers.

There are no ring outliers.

No monomer is involved in short contacts.

##### 5.7 Other polymers (i)

There are no such residues in this entry.

PRELIMINARY VALIDATION REPORT

#### 5.8 Polymer linkage issues (i)

There are no chain breaks in this entry.

PRELIMINARY VALIDATION REPORT

### 6 Fit of model and data (i)

#### 6.1 Protein, DNA and RNA chains (i)

In the following table, the column labelled '#RSRZ> 2' contains the number (and percentage) of RSRZ outliers, followed by percent RSRZ outliers for the chain as percentile scores relative to all X-ray entries and entries of similar resolution. The OWAB column contains the minimum, median, $95^{th}$ percentile and maximum values of the occupancy-weighted average B-factor per residue. The column labelled 'Q< 0.9' lists the number of (and percentage) of residues with an average occupancy less than 0.9.

| Mol | Chain | Analysed | <RSRZ> | #RSRZ>2 | | | OWAB(Å$^2$) | Q<0.9 |
|---|---|---|---|---|---|---|---|---|
| 1 | A | 151/151 (100%) | -0.48 | 0 | 100 | 100 | 36, 51, 64, 80 | 0 |
| 1 | G | 151/151 (100%) | -0.39 | 0 | 100 | 100 | 38, 52, 69, 75 | 0 |
| 2 | B | 94/94 (100%) | -0.15 | 1 (1%) | 80 | 77 | 34, 47, 66, 74 | 0 |
| 2 | H | 94/94 (100%) | -0.24 | 0 | 100 | 100 | 35, 49, 61, 77 | 0 |
| 3 | E | 4/4 (100%) | -0.09 | 0 | 100 | 100 | 49, 49, 53, 59 | 0 |
| 3 | I | 4/4 (100%) | -0.39 | 0 | 100 | 100 | 44, 48, 53, 56 | 0 |
| 3 | L | 4/4 (100%) | 0.87 | 0 | 100 | 100 | 74, 76, 84, 88 | 0 |
| 4 | C | 145/145 (100%) | -0.36 | 0 | 100 | 100 | 40, 55, 67, 86 | 0 |
| 5 | D | 94/95 (98%) | -0.20 | 1 (1%) | 80 | 77 | 34, 48, 75, 82 | 0 |
| 6 | J | 144/144 (100%) | -0.11 | 1 (0%) | 87 | 85 | 44, 65, 77, 86 | 0 |
| 7 | K | 88/96 (91%) | -0.14 | 1 (1%) | 80 | 77 | 41, 56, 84, 93 | 0 |
| All | All | 973/982 (99%) | -0.27 | 4 (0%) | 92 | 91 | 34, 54, 73, 93 | 0 |

All (4) RSRZ outliers are listed below:

| Mol | Chain | Res | Type | RSRZ |
|---|---|---|---|---|
| 5 | D | 197 | VAL | 4.8 |
| 6 | J | 61 | LEU | 2.1 |
| 2 | B | 265 | PRO | 2.0 |
| 7 | K | 271 | GLY | 2.0 |

#### 6.2 Non-standard residues in protein, DNA, RNA chains (i)

There are no non-standard protein/DNA/RNA residues in this entry.

PRELIMINARY VALIDATION REPORT

#### 6.3 Carbohydrates (i)

There are no carbohydrates in this entry.

#### 6.4 Ligands (i)

In the following table, the Atoms column lists the number of modelled atoms in the group and the number defined in the chemical component dictionary. The B-factors column lists the minimum, median, $95^{th}$ percentile and maximum values of B factors of atoms in the group. The column labelled 'Q< 0.9' lists the number of atoms with occupancy less than 0.9.

| Mol | Type | Chain | Res | Atoms | RSCC | RSR | B-factors(Å$^2$) | Q<0.9 |
|---|---|---|---|---|---|---|---|---|
| 8 | VAL | F | 301 | 7/? | 0.90 | 0.16 | 60,67,68,70 | 0 |

#### 6.5 Other polymers (i)

There are no such residues in this entry.

PRELIMINARY VALIDATION REPORT
